# Genetic, morphological, and niche variation in the widely hybridizing *Rhus integrifolia-Rhus ovata* species complex

**DOI:** 10.1101/2020.04.30.070813

**Authors:** Craig F. Barrett, Joshua Lambert, Mathilda V. Santee, Brandon T. Sinn, Samuel V. Skibicki, Heather M. Stephens, Hana Thixton

## Abstract

Hybridization and introgression are common processes among numerous plant species that present both challenges and opportunities for studies of species delimitation, phylogenetics, taxonomy, and adaptation. *Rhus integrifolia* and *R. ovata* are two ecologically important shrubs native to the southwestern USA and Mexico, and are known to hybridize frequently, but the morphological, genetic, and ecological implications of hybridization in these species are poorly studied on a broad geographic scale. Analyses were conducted using leaf morphology, genetic variation of plastid and nuclear loci, and species distribution models for both species and their putative hybrid introgressants across 19 localities in California and Arizona, USA. These analyses revealed evidence for morphological and genetic distinction among localities comprising putative parental species, but a high degree of morpho-genetic intermediacy among localities with putative hybrids. Comparison of morphological and genetic population structure among localities revealed evidence for putative local adaptation or widespread phenotypic plasticity. Multiple regression models identified a weak but statistically significant negative association between leaf area and precipitation. Finally, species distribution modeling inferred northward range shifts over time, with both species predicted to occupy more coastal regions in the future, possibly increasing the frequency of hybridization among them. These findings underscore the importance of integrative assessment of multiple data sources in the study of hybridizing species and highlight the *Rhus integrifolia-ovata* complex as a powerful model for investigating the adaptive implications of hybridization.

## 1 INTRODUCTION

Hybridization is a hallmark of many plant species complexes, often blurring species boundaries (Stebbins, 1969; Rieseberg and Soltis, 1991; Petit and Excoffier, 2009). The interplay between divergent selection among parental species in terms of maximizing reproductive success in a particular niche vs. the frequency and extent of gene flow remains a key issue in evolutionary biology (Slatkin, 1987; Burke et al., 1998; Rieseberg et al., 1999; Saccheri and Hanski, 2006; Sork et al., 2016). The potential outcomes of hybrid introgression are diverse and context-dependent, including: 1) lower fitness among hybrid offspring reinforcing species boundaries among distinct parental species (e.g. Rieseberg et al., 1999; Hoskin et al., 2005); 2) higher fitness among offspring in novel or intermediate environments relative to parental species allowing invasion of marginal or novel niches, ultimately leading to ecological specialization and eventual speciation (e.g. Rieseberg et al., 1995; Ellstrand, 2003; Mavarez et al., 2006; Soltis and Soltis, 2009; Abbott et al., 2013); 3) little to no fitness consequences among offspring, with introgression simply serving as a vehicle for ‘neutral’ genetic exchange among parental species (e.g. Gavrilets and Cruzan, 1998); 4) the exchange of novel, adaptive genetic variation via gene flow between parental species through introgressants (Ellstrand and Schierenbeck, 2000; Arnold, 2004; Hegarty and Hiscock, 2004).

A ‘classic’ example of hybrid introgression is that of *R. integrifolia* (Nutt.) Benth. & Hook. f. ex Rothr. and *Rhus ovata* S. Watson, two ecologically important shrubs native to southwestern North America (Barkley, 1937; Young, 1974). Both are major structural components of coastal scrub and chaparral ecosystems, respectively, and are important in erosion control, as native ornamental shrubs/trees, and as a source of food and shelter for wildlife. *Rhus integrifolia* occupies coastal scrub habitats in California (USA), Baja California (Mexico), and outlying islands. *Rhus ovata* occurs in coastal mountains in the chaparral regions of California and northwestern Mexico and is also disjunct to interior chaparral habitats of central Arizona (USA), separated from Californian conspecifics by the Sonoran and Mojave deserts (Montalvo et al., 2017). Both species are gynodioecious, with hermaphroditic and male-sterile individuals frequently occurring in the same populations. This reproductive strategy has evolved numerous times in plants and is hypothesized to promote outcrossing and thereby reduce the deleterious effects of inbreeding depression (Barkley, 1937; Munz and Keck, 1959; Young, 1972; 1974; Freeman et al., 1997; Barrett, 2002).

*Rhus integrifolia* has relatively small, flat, often toothed, obovate-obelliptic leaves (3-6 cm in length). *Rhus ovata* has broad, waxy, ovate-deltoid leaves that fold along the midrib of the abaxial surface into a characteristic “taco” shape, which is likely an adaptation to hot, arid summers in mid-lower montane chaparral zones (leaves 4-11 cm in length). These morphologies may represent extremes on an environmental continuum based on proximity to the Pacific Ocean, moisture, and temperature fluctuations in a diverse, heterogeneous range from coastal California and Baja California to interior Arizona (Young, 1974; Montalvo et al., 2017). Californian populations of these two species show varying degrees of morphological intermediacy due to introgression at intermediate elevations (Figs. 1; S1), where they are often sympatric; i.e. in regions where the mountains abruptly meet the coast (Barkley, 1937; Young, 1974). Arizonan populations of *R. ovata*, on the other hand, are allopatrically separated from *R*. *integrifolia* or any putatively introgressant populations of *R. ovata*, and thus may represent a “pure” form of *R. ovata*.

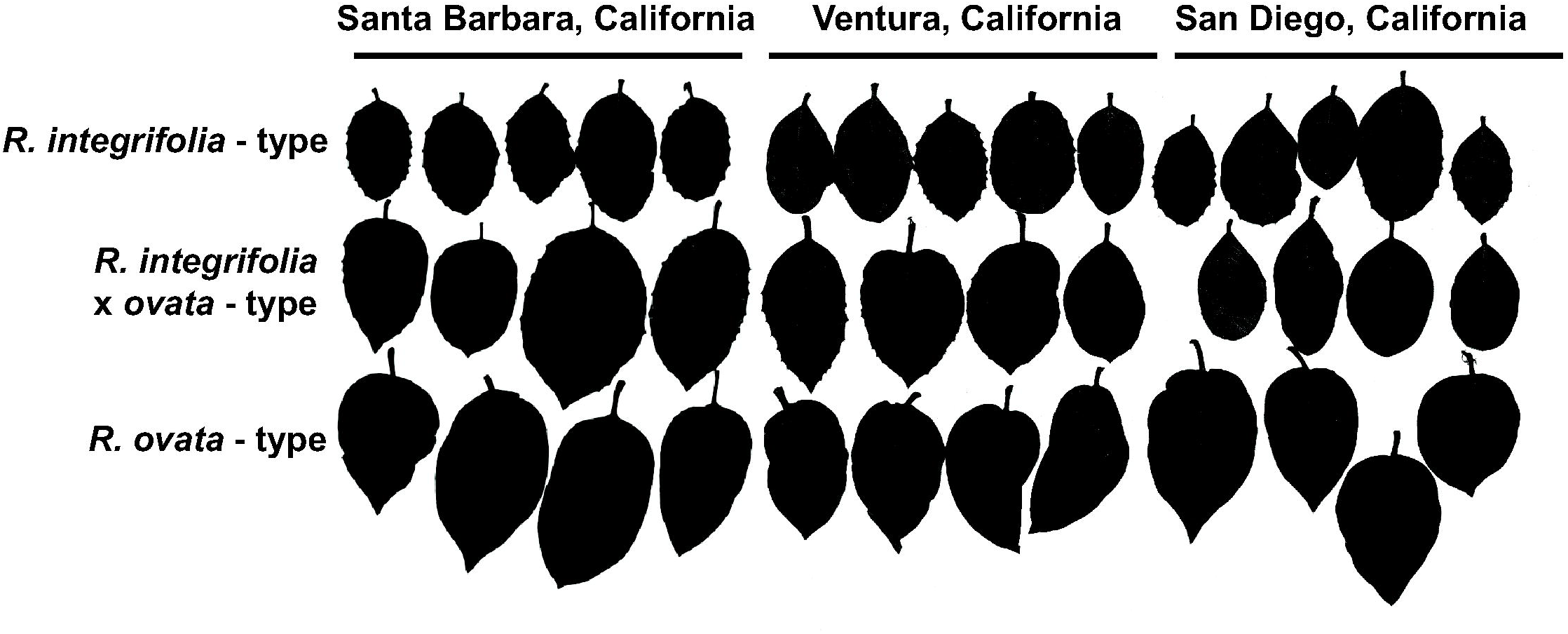

The two species are estimated to have diverged ca. 3 million years ago (mya) +/- 1.6 mya (Miller et al., 2001, Yi et al., 2004). Fossils attributed to both species have been found at inland sites in Nevada, farther north than the current distribution of either species, dating back to the Miocene and even Pliocene (Young, 1974). Thus, these two species may have undergone several periods of contracting and expanding distributions, being both allopatric and sympatric over hundreds of thousands to a few million years, e.g. spanning several of the Pleistocene glaciations.

Young (1972; 1974) conducted meticulous studies of the breeding systems and patterns of introgressive hybridization in these two species, based on samples from two “pure” localities of each species and one sympatric locality, demonstrating intermediacy in leaf and floral traits in the sympatric population. However, Young (1974) conceded that the two “pure” populations of *R. ovata* displayed some intermediate features akin to *R. integrifolia*, and could not rule out that these populations may be the result of either ‘ancient’ introgression, or introgression in the immediate past followed by backcrossing with more ‘pure’ forms of *R. ovata*. Further, Young (1974) found limited evidence for clinal variation in leaf length and width associated with latitude within *R. ovata*, with shorter, more narrow leaves in the southernmost population compared to larger, broader leaves in the northern population, though this finding is based on comparison of only two populations.

Despite these earlier studies, a quantitative assessment of range-wide variation in morphology, genetic diversity, and abiotic niche requirements is lacking for these two ecologically important species. Here we use plastid and nuclear DNA sequences, leaf morphometrics, and species distribution models to characterize patterns of differentiation and hybrid introgression in the *Rhus integrifola*-*ovata* complex, addressing the following questions: 1) What is the extent of leaf morphological variation across the geographic ranges of *R. ovata* and *R. integrifolia*, and within introgressed populations, and how to these relate to environmental variation? 2) Do plastid and nuclear DNA show distinct patterns of population structuring across the allopatric and sympatric portions of their ranges, and what is the extent of genetic evidence for introgression? Specifically, are the disjunct Californian and Arizonan populations genetically distinct or do they display evidence of current or historical introgression (shared haplotypes) with populations in the sympatric range? 3) How do the inferred environmental niches of *R. integrifolia* and *R. ovata* differ in the present, past, and future, and what is their degree of niche overlap?

## 2 MATERIALS AND METHODS

### 2.1 Sampling

Leaf material and voucher specimens were sampled from 19 localities across California and Arizona (Table 1). Localities correspond to putatively “pure” populations of *R. ovata* (n = 12), *R. integrifolia* (n = 4), or of ‘mixed’ populations with signs of morphological intermediacy (*R. integrifolia* × *ovata*; n = 3; Fig. 1). No populations were included from the Channel Islands or Baja California due to logistical challenges of collecting material.

**TABLE 1.** Collection locality information for *R. integrifolia, R. ovata*, and *R. integrifolia* × *ovata*. US state codes: CA = California, AZ = Arizona.

| Species present | Locality Code | Locality | County, US state | Latitude, Longitude | Elevation (m) |
| --- | --- | --- | --- | --- | --- |
| *R. ovata* | GIAZ-GL | Globe, Route 77/60, Tonto National Forest | Gila, AZ | 33.552416, -110.677328 | 1330 |
| *R. ovata* | GIAZ-6S | Six-shooter Canyon, Tonto National Forest | Gila, AZ | 33.339374, -110.784457 | 1218 |
| *R. ovata* | GIAZ-KC | Kellner Canyon Road, Tonto National Forest | Gila, AZ | 33.334900, -110.830900 | 1343 |
| *R. integrifolia* | LACA-PV | Palos Verdes Peninsula | Los Angeles, CA | 33.743931, -118.411408 | 47 |
| *R. ovata* | LACA-CF | Chantry Flat, Angeles National Forest | Los Angeles, CA | 34.190420, -118.022213 | 605 |
| *R. ovata* | LACA-LH | Lake Hughes, Angeles National Forest | Los Angeles, CA | 34.560400, -118.491109 | 551 |
| *R. ovata* | SDCA-BG | Banner Grade, Route 78 | San Diego, CA | 33.084277, -116.563568 | 307 |
| *R. integrifolia* × *ovata* | SDCA-SC | Sycamore Canyon Road | San Diego, CA | 32.942893, -116.985358 | 289 |
| *R. ovata* | SDCA-PM | Palomar Mountain, South Grade, Cleveland National Forest | San Diego, CA | 33.305918, -116.871002 | 1401 |
| *R. integrifolia* | SDCA-EF | Elfin Forest Road | San Diego, CA | 33.086753, -117.147970 | 150 |
| *R. ovata* | SDCA-RG | Rainbow Glen Road | San Diego, CA | 33.415497, -117.174885 | 402 |
| *R. ovata* | SBCA-CS | Cold Spring Trail, Los Padres National Forest | Santa Barbara, CA | 34.458688, -119.648858 | 394 |
| *R. ovata* | SBCA-ST | Snyder Trail, Los Padres National Forest | Santa Barbara, CA | 34.536375, -119.789871 | 520 |
| *R. integrifolia* | SBCA-EC | El Capitan State Beach | Santa Barbara, CA | 34.461526, -120.012142 | 12 |
| *R. integrifolia* | VECA-BB | Beach to Backcountry Trail, Gaviota State Park | Santa Barbara, CA | 34.482077, -120.236825 | 133 |
| *R. integrifolia* × *ovata* | SBCA-BG | Los Padres National Forest, near Santa Barbara Botanical Garden | Santa Barbara, CA | 34.467932, -119.708093 | 227 |
| *R. integrifolia* × *ovata* | VECA-LJ | La Jolla Canyon, Los Padres National Forest | Ventura, CA | 34.092930, -119.039087 | 206 |
| *R. ovata* | YAAZ-PR | Route 89, Prescott National Forest, near Prescott | Yavapai, AZ | 34.423816, -112.553975 | 1680 |
| *R. ovata* | YAAZ-BA | Route 96, Baghdad, AZ | Yavapai, AZ | 34.557514, -113.160571 | 1064 |

### 2.2 Morphology

Leaves were collected from the basal-most position along a branch, in order to reduce the effect of intra-individual morphological heterogeneity. We were unable to include floral/fruit characters for every locality and instead focused on leaf characters exclusively, which have been shown to be useful indicators of introgression between *R. integrifolia* and *R. ovata* (Young, 1974). Pressed leaves were scanned at 1200 DPI on a Brother TN630 scanner along with a flat 6cm ruler (Fisher Scientific, Waltham, Massachusetts, USA). The software ImageJ2 (Rueden et al., 2017) was used to measure five continuous characters: lamina length, lamina width at widest point, lamina width at 1/4 distance from apex, lamina width at 1/4 distance from base, and petiole length. We scored two meristic characters: number of secondary veins along the adaxial lamina surface on the left side, and the total number of teeth along the lamina margin. We scored seven discrete characters: teeth (present/absent); acute lamina apex (present/absent); base of lamina (acute/truncate/cordate); lamina folding (folded/flat/wavy); red lamina margin (present/absent); basal lamina lobing (present/absent); and lamina shape (ovate-deltoid/oval-elliptic/obovate-obelliptic).

We conducted Principal Components Analysis (PCA) using a correlation matrix in PAST v.3 (Hammer et al., 2001) on all characters and used the ‘biplot’ function to investigate the relative contributions of each character to each resulting PC axis. We used the scree plot and ‘broken stick’ method in PAST to assess the number of PC axes contributing significantly to total variation. We plotted the density of individual scores along PC1 in R using the violin and jitter plot functions in the R package ‘ggplot2’ (Wickham, 2016). We interpreted scores and density of individuals along PC1 as a proxy of a morphological hybrid index among four a priori groupings of taxa/localities: coastal and interior Californian *Rhus ovata*; coastal Californian *R. integrifolia*; interior Arizonan *R. ovata*, and localities with *R. ovata, R. integrifolia*, and their introgressants (*R. integrifolia* × *ovata*). We conducted a two-way nonparametric multivariate analysis of variance with Gower transformation for ‘mixed’ data (NP-MANOVA) to evaluate statistically significant multivariate differences in leaf morphology among groupings in PAST v.3, partitioning among two levels: grouping and locality within grouping.

### 2.3 Leaf area and environmental variation

We constructed multiple regression models in order to investigate the relationship of leaf area to environmental variation. We analyzed the log_10_ of leaf area, quantified using ImageJ2, in association with four composite environmental predictor variables. We did not calculate specific leaf area (i.e. leaf area/leaf dry mass) because we were unable to dry the leaves simultaneously under identical conditions, which could have been a source of bias. Nineteen BIOCLIM environmental variables were downloaded for each sampling locality at 2.5 arc-minute resolution from https://www.worldclim.org using the R package ‘raster’ (Hijmans, 2019). Then, PCA was conducted on the nineteen BIOCLIM variables using a correlation matrix in PASTv.3.

Temperature and precipitation-related variables were analyzed separately in order to investigate their effects individually. A broken-stick analysis was conducted as above. The first two PCs for temperature and precipitation were retained for downstream regression analyses. In addition, a binary grouping variable was included each for localities containing *R. ovata* (CA), *R. ovata* (AZ), *R. integrifolia* (CA) and *R. integrifolia* × *ovata* (CA) (n = 4 groups), to account for variation in leaf area among groups.

We included all combinations of PC1 (temperature), PC2 (temperature), PC1 (precipitation), PC2 (precipitation), and ‘group’ in the models. Relative importance of each term (environmental PCs, group, and interaction terms) was assessed via significance in the models. The full model was specified as: log_10_ leaf area ∼ PC1_temp_ + PC2_temp_ + PC1_precip_ + PC2_precip_ + group + interaction terms + error. Interaction terms included all pairwise combinations of temperature × precipitation PCs, and of each PC × group. Loadings scores for each BIOCLIM variable with coefficients above a threshold of 0.25 were interpreted as the variables most strongly associated with each PC. We ran eight nested models, dropping the *R. integrifolia* (CA) grouping variable for those models that included a group effect. All analyses were conducted in R using the ‘lm’ function. Tables were summarized with ‘jtools’ (Long, 2019) and ‘huxtable’ (Hugh-Jones, 2018) in R, reporting R^2^ and R^2^-adjusted values for each model, as well as regression coefficients, standard errors, and significance for each predictor and interaction term.

### 2.4 Plastid and nuclear ITS variation

Approximately 1 cm^2^ of leaf material was removed from the center of each leaf adjacent to the midrib prior to pressing as not to obscure morphological features of the leaf, and DNA was extracted following a modified CTAB protocol (Doyle and Doyle, 1987), at 1/5 volume. Total genomic DNA was subjected to PCR amplification of one nuclear and two plastid regions. We amplified the nuclear internal transcribed spacer (ITS) with primers ITS1 and ITS4 (White et al., 1990), the plastid *ndhC-trnV*^*UAC*^ spacer with primers rhus-ndhC-F (5’ AGCAGAAACATAGACGAACTCTCC 3’) and rhus-trnV-R (5’ GTCTACGGTTCGAGTCCGTATAGC 3’), and the plastid *rpl16-rps3* intergenic spacer with primers rhus-rpl16-F (5’ GGTTCCATCGTTCCCATTGCTTCT 3’) and rhus-rps3-R (5’ TGTAGCCGCAGAATAATAAGACT 3’). These regions were chosen based on a previous assessment of high-variation plastid markers based on complete plastid genomes of *R. integrifolia* and *R. ovata* (NCBI GenBank accession numbers MT024991-MT024993; Barrett, in review). Reactions were carried out in 25μl volumes, with 12.5μl Apex PCR Master Mix (Genesee Scientific, San Diego, California, USA), nine μl pure water, 0.2μM of each primer, 0.5μl 5M Betaine, and 20-100ng template DNA in one μl Tris-EDTA Buffer (pH = 8.0). PCR conditions for ITS consisted of 95°C for 3 min, 30 cycles of 95°C (30 sec), 55°C (45 sec), and 72°C (90 sec), with a final extension of 72°C for 10 min. Conditions for plastid loci differed only by the annealing step (60°C for 30 sec). PCR products were visualized on 1% agarose gels and cleaned with 1.8x volume of AxyPrep FragmentSelect magnetic beads (Corning-Axygen, Corning, New York, USA), followed by two washes with 80% ethanol. PCR products were cleaned with Sephadex G-50 fine medium (70g/L; GE Healthcare, Chicago, Illinois, USA), centrifuged through a 96-well filter plate (Phenix Research, Accident, Maryland, USA), quantified via NanoDrop spectrophotometry (ThermoFisher), and diluted to 30ng/μl. PCR products were sequenced on both strands using the same primers as for amplification, following manufacturer protocols (Applied Biosystems BigDye v.3.1 cycle sequencing kit, Life Technologies, Waltham, Massachusetts, USA) on an ABI 3130XL Genetic Analyzer at the West Virginia University Genomics Core Facility.

Resulting chromatograms were edited in Geneious R10 (http://www.geneious.com) and consensus sequences were aligned with MAFFT v.7 (Katoh and Standley, 2013) under default parameters (gap opening penalty = 2.0, offset value = 0.5). For ITS electropherograms, all bi- allelic, heterozygous sites were manually checked in Geneious and scored using IUPAC ambiguity codes. Alignments for each locus were conducted with MAFFT (gap opening = 3, offset = 0.5), adjusted manually at the margins in Geneious, and trimmed to remove ambiguous calls near the priming sites. SeqPhase (Flot, 2010) and PHASE (Stephens et al., 2001) were used to determine ITS alleles, with a 90% posterior probability per heterozygous site, using sampled homozygous sequences as prior information. A variable minisatellite repeat in the *ndhC-trnV* spacer was coded as a single, multistate character (ATT TTT TT[K] ATT ATT AAT TAT T). Plastid loci were concatenated and analyzed as a single alignment. Sequences are deposited in NCBI GenBank (Accession numbers: XXXXXXX-XXXXXXX), and alignment/morphological data deposited in Dryad (XXXXXX).

Haplotype networks were constructed in PopART v.1.7 (Leigh and Bryant, 2015), using the ‘TCS network’ option (Clement et al., 2002) with a 95% connection limit. Locality codes and GPS information (decimal degrees) were then added to a NEXUS file of the plastid and ITS alignments to map haplotype frequencies using PopART and edited in Adobe Illustrator v. 24.1.1 (Adobe Inc., 2019).

### 2.5 Population genetics

Population genetic analyses were carried out in ARELQUIN v.3.5 (Excoffier and Lischer, 2010) for both plastid and nuclear datasets. Analysis of molecular variance (AMOVA; Excoffier et al., 1992) was conducted for each dataset and partitioned by locality and grouping (as in Table 1) to quantify the proportion of variation at different hierarchical levels, and to assess the degree of population structuring across the range of this complex. Pairwise comparisons of Φ_ST_ and their significance were further conducted among localities in ARLEQUIN. The values N_ST_ and G_ST_ were compared in SPADS v.1.0 (Dellicour and Mardulyn, 2014) in order to further test significance of population genetic structure among localities. Population structure was deemed to be statistically significant if N_ST_, which accounts for nucleotide sequence divergence, was significantly greater than G_ST_, which treats alleles as discrete units.

We conducted an overall comparison of the relative degrees of morphological and genetic variation among localities using the ‘Pstat’ package in R (da Silva and da Silva, 2018). This package calculates ‘P_ST_’, an analog of ‘Q_ST_’, which may serve as a proxy for genetically determined morphological variation over a range of levels of additive genetic variation and narrow-sense heritability. We used individual scores along Reist-transformed (Reist, 1985) morphological PCs 1 and 2 as metrics of multivariate morphological variation (see above), and global estimates of Φ_ST_ from plastid and nuclear ITS data, respectively. Without information from common garden and reciprocal transplant experiments (which would have been infeasible for the current study), P_ST_ > Φ_ST_ may indicate localized adaptive evolution via divergent selection among populations but may also reflect plastic responses to environmental variation (Brommer, 2011). Given the lack of studies on genetically determined morphological variation in both species of *Rhus* and their introgressant populations, we used this as an exploratory tool to distinguish a situation in which measures of the degree of morphological variation (P_ST_) outrank the degree of neutral genetic differentiation (Φ_ST_) without distinguishing between genetic and environmental factors, but instead tested this relationship over a range of potential heritability.

### 2.6 Species distribution modeling

Occurrence data were downloaded from the Global Biodiversity Information Facility (GBIF) using the ‘rgbif’ package for R. The R package ‘CoordinateCleaner’ (Zizka et al., 2018) was used to filter occurrence data on several criteria, including latitude (below 35°N, corresponding to the northern limits the native ranges of both species), preserved specimens only, year (post- 1930), occurrences missing coordinates, coordinate uncertainty, and range outliers.

We used the R packages ‘raster’ (Hijmans et al., 2019) and ‘sp’ (Pebesma and Bivand, 2005) to procure 19 BIOCLIM variables from the WorldClim database (Fick and Hijmans, 2017), corresponding to the cleaned *R. integrifolia* and *R. ovata* datasets based on their GPS coordinates (10 km resolution), with the addition of the 19 sampled localities from this study. We log_10_- transformed the BIOCLIM data after adjusting negative and zero values (by adding 100 to these variables). We subjected the dataset to Principal Components analysis via a correlation matrix in PAST v.3, and plotted *R. integrifolia* and *R. ovata* occurrence data with the addition of new collections. We further used NP-MANOVA to test for significant multivariate differences in environmental variables between *R. integrifolia* and *R. ovata* in PAST.

Species distribution models (SDM) were inferred using MaxEnt (version 3.4.1; Phillips et al., 2006; 2017) via the R package Dismo. Last Glacial Maximum (∼22 kya), mid-Holocene (∼6 kya), contemporary, and future (IPPC5 2070) bioclimatic predictor variable layers were obtained via WorldClim (worldclim.org) at 2.5 arc-minute resolution. Last Glacial Maximum and mid-Holocene predictor layers were generated using the fourth release of the Community Climate System Model (Gent et al., 2010). We assessed the impact of future climate change on species distributions by inferring future habitable ranges using both the 2.6 and 8.5 greenhouse gas representative concentration pathway (RCP) scenarios. Climate layer inclusion was filtered using Pearson correlation thresholds of 0.85 and -0.85. Analyses of niche equivalency were conducted per Barrett et al. (2019).

## 3 RESULTS

### 3.1 Morphology

Principal Components Analysis of 14 morphological characters revealed clear separation among individuals from localities hypothesized to represent “pure” forms of *R. integrifolia* and *R. ovata* in CA and AZ, as indicated by non-overlapping 95% Cis of PC axes 1 and 2 (Fig. 2A). Individuals from localities with hypothesized introgression between the two species overlapped broadly in multivariate space with both species. Among localities with *R. ovata*-type morphologies only, the CA and AZ individuals showed overlap, but the AZ populations only occupied a relatively small region of multivariate space, with AZ individuals contained within the 95% CI of the CA individuals. PC1 explained 48.99% of the total variation in morphological features, while PC2 explained 10.63% (Table 2; S3). PC1 was largely determined by lamina length/width characters, petiole length, the number of secondary veins, lamina apex shape, leaf folding pattern, and overall leaf shape (Figs. 2B; S3). PC2 was largely determined leaf shape characters (overall shape, folding, basal lamina shape), and the pattern of teeth on the lamina margin.

**Table 2.**
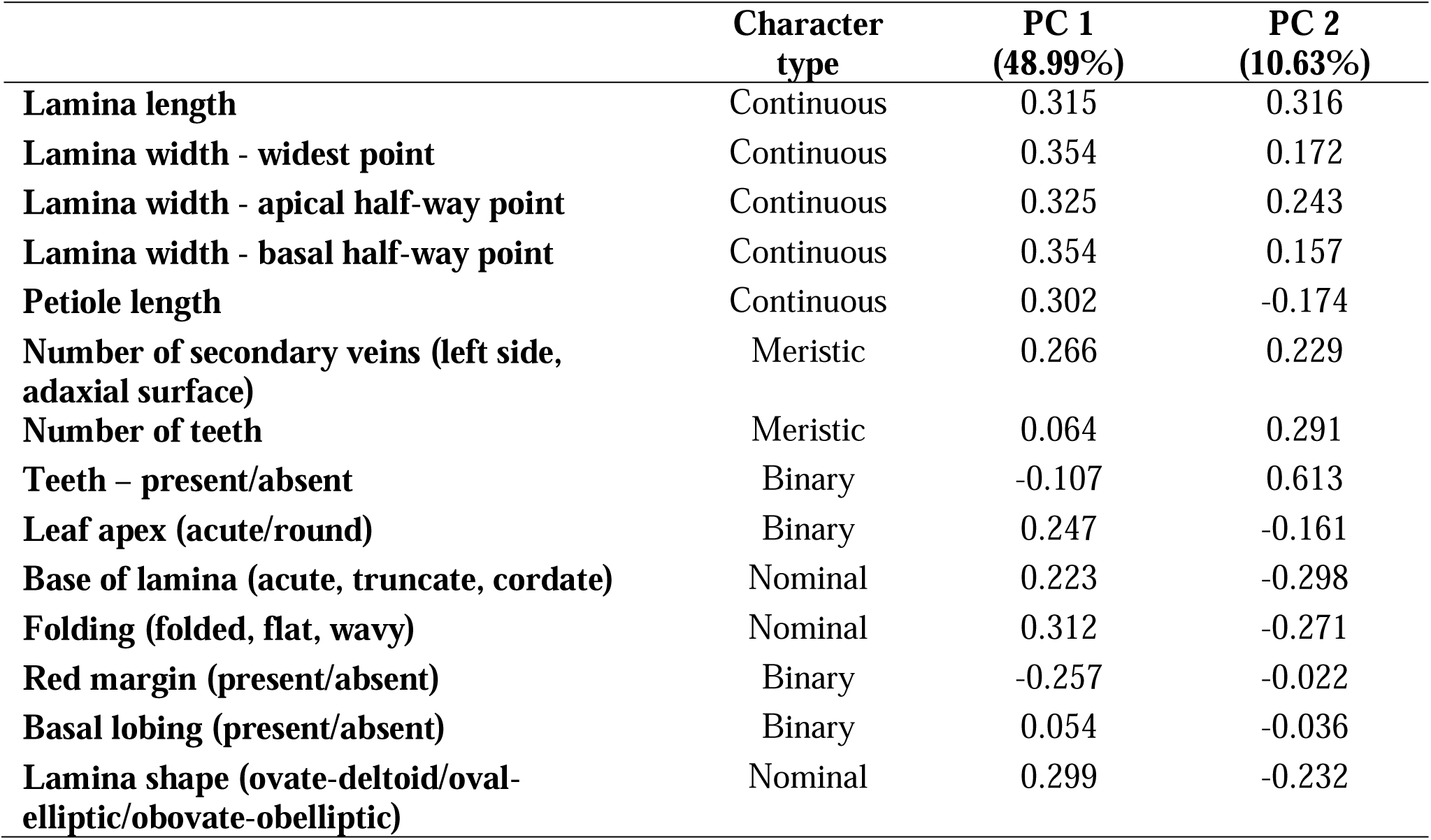
Morphological character definitions and PCA loadings (PCs 1-2) based on a correlation matrix in PASTv.3.

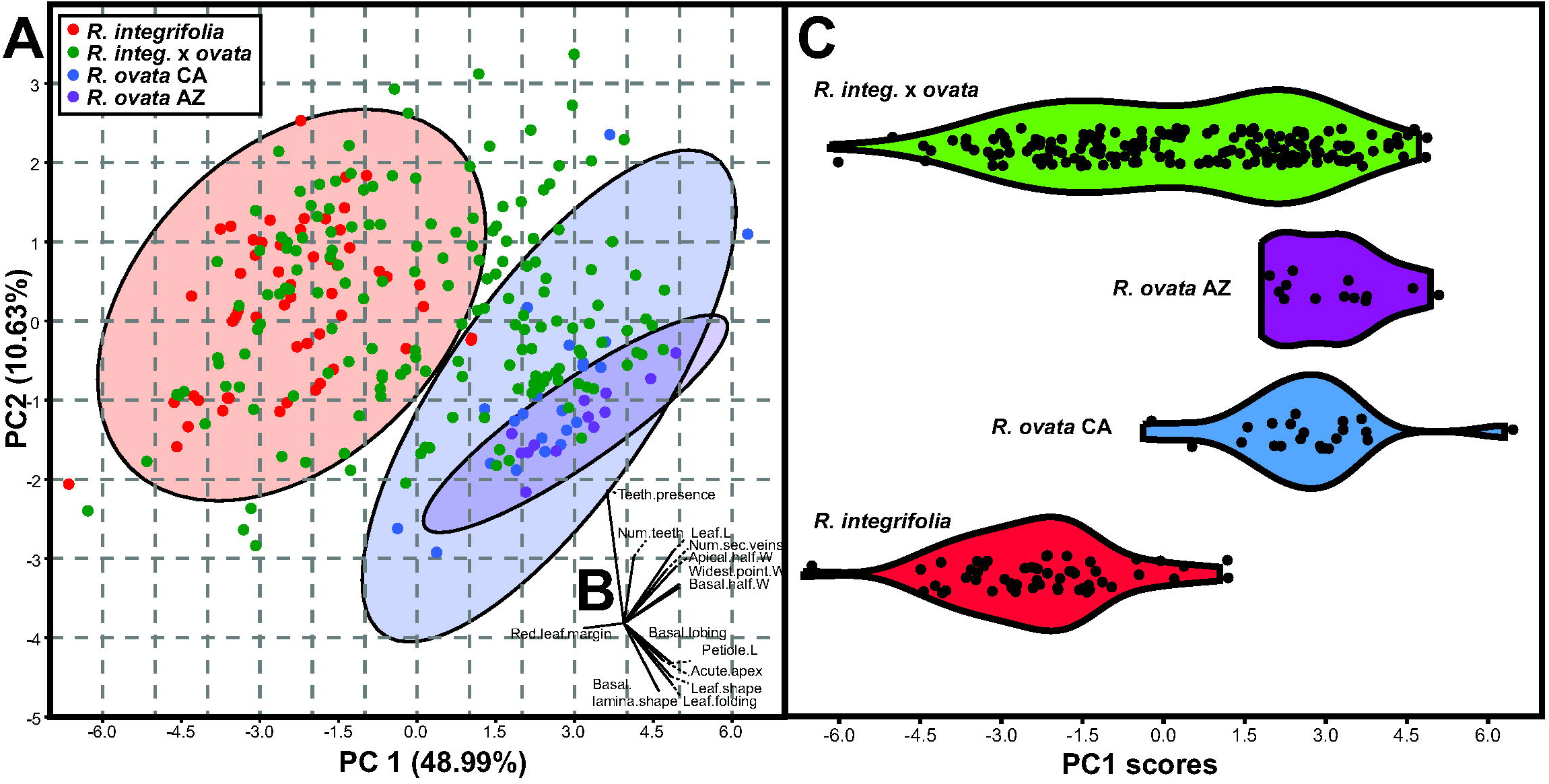

Morphological variation is further illustrated by a violin plot of PC1 for *R. integrifolia* vs. *R. ovata* (both CA and AZ localities), which showed clear separation among species, with individuals from introgressant populations covering nearly the entire PC1-ranges of both (Fig. 2C). A two-way NP-MANOVA of Gower-transformed data suggested that the groupings from Fig. 2A differed significantly in multivariate leaf morphological space (Table S4., F = 9.897, p = 0.0001), and also that individuals from different sampling localities differed significantly within each grouping (F = 3.511, p = 0.0001), with no significant interaction among grouping and locality factors (F = -11.970, p = 1.0).

### 3.2 Leaf area and environmental variation

Principal components analyses of 11 BIOCLIM temperature variables and eight precipitation variables indicated that the first two temperature and precipitation PCs were significant, based on a broken stick analysis (Fig. S5). Based on loading scores from the PCA: PC1_temp_ reflected overall temperature magnitude (60.6% of total variance); PC2_temp_ reflected temperature variation (31,1% of total variance, e.g. isothermality, mean diurnal range, annual temperature range); PC1_precip_ reflected overall precipitation magnitude (62.7% of total variance); and PC2_temp_ reflected variation in precipitation (24.3% of total variance, e.g. precipitation seasonality) (Fig. S5).

Multiple regression models for log_10_ leaf area and environmental variables are summarized in Table 3. Model 1 included only the four environmental PCs and indicated a significant negative relationship between PC1_precip_ and leaf area (coefficient = -0.04, standard error = 0.01, p < 0.01; reported hereafter as ‘coef’, ‘se, and ‘p,’ respectively). Model 2 included only the binary grouping variables and indicated that the ‘groups’ each captured a significant amount of variation in leaf area (coef = 0.20, se = 0.03; coef = 0.30, se = 0.04; coef = 0.33, se = 0.05; p < 0.001 for all). Model 3 included environmental PCs and group, further indicating that groups captured most of the variation in leaf area. Model 4 included environmental PCs and all pairwise interaction terms between temperature and precipitation PCs, and indicated a significant negative association between leaf area and PC1_temp_ (coef = -0.10, se = 0.02, p < 0.001); a significant positive association with PC2_temp_ (coef = 0.18, se = 0.04, p < 0.001), and a significant negative relationship with PC1_precip_. (coef = -0.06; se = 0.02, p < 0.01). This model also indicated a significant interaction among PC1_precip_ and both temperature PCs (coef = -0.03, se = 0.01, p < 0.001 for PC1_temp_; and coef = 0.02, se = 0.00, p < 0.001 for PC2_temp_, respectively).

**Table 3.** Multiple regression model summary for leaf area vs. PCs 1 and 2 (temperature) and PCs 1 and 2 (precipitation). * p < 0.05, ** p < 0.01, *** p < 0.001. See text for further explanation of the models. Model 8 is not shown, as none of the terms were significant (including all terms and their interactions).

|  | **Model 1** |  | **Model 2** |  | **Model 3** |  | **Model 4** |  | **Model 5** |  | **Model 6** |  | **Model 7** |  |
| --- | --- | --- | --- | --- | --- | --- | --- | --- | --- | --- | --- | --- | --- | --- |
|  | Coef. | SE | Coef. | SE | Coef. | SE | Coef. | SE | Coef. | SE | Coef. | SE | Coef. | SE |
| (Intercept) | 5.91 *** | 0.01 | 5.74 *** | 0.02 | 5.75 *** | 0.04 | 6.00 *** | 0.04 | 5.84 *** | 0.06 | 5.47 *** | 0.56 | 5.14 *** | 0.44 |
| PC1.temp | 0.00 | 0.01 |  |  | -0.01 | 0.01 | **-0.10** *** | 0.02 | -0.06 | 0.03 | 0.18 | 0.32 | -0.01 | 0.02 |
| PC2.temp | -0.02 | 0.01 |  |  | -0.00 | 0.02 | **0.18** *** | 0.04 | 0.09 | 0.05 | -0.39 | 0.58 | -0.03 | 0.03 |
| PC1.precip | **-0.04** ** | 0.01 |  |  | -0.00 | 0.02 | **-0.06** ** | 0.02 | -0.02 | 0.03 | **-0.08** * | 0.03 | 0.42 | 0.32 |
| PC2.precip | 0.01 | 0.01 |  |  | -0.02 | 0.01 | -0.05 | 0.03 | -0.04 | 0.04 | -0.04 | 0.02 | -0.36 | 0.21 |
| group [2] *R. integ.* × *ovata* |  |  | **0.20** *** | 0.03 | **0.20** *** | 0.03 |  |  | **0.13** ** | 0.04 | 0.65 | 0.58 | 0.78 | 0.43 |
| group [3] *R. ovata* (CA) |  |  | **0.30** *** | 0.04 | **0.32** *** | 0.06 |  |  | **0.22** ** | 0.07 | 0.50 | 0.56 | 0.84 | 0.44 |
| group [4] *R. ovata* (AZ) |  |  | **0.33** *** | 0.05 | 0.25 | 0.25 |  |  | 0.50 | 0.28 | -0.13 | 0.60 | **1.43** * | 0.65 |
| PC1.temp * PC1.precip |  |  |  |  |  |  | **-0.03** *** | 0.01 | **-0.02** * | 0.01 |  |  |  |  |
| PC1.temp * PC2.precip |  |  |  |  |  |  | -0.01 | 0.01 | -0.01 | 0.01 |  |  |  |  |
| PC2.temp * PC1.precip |  |  |  |  |  |  | **0.02** *** | 0.00 | **0.01** ** | 0.01 |  |  |  |  |
| PC2.temp * PC2.precip |  |  |  |  |  |  | -0.01 | 0.01 | -0.00 | 0.01 |  |  |  |  |
| PC1.temp * group [2] |  |  |  |  |  |  |  |  |  |  | -0.31 | 0.34 |  |  |
| PC1.temp * group [3] |  |  |  |  |  |  |  |  |  |  | -0.17 | 0.32 |  |  |
| PC1.temp * group [4] |  |  |  |  |  |  |  |  |  |  | -0.23 | 0.32 |  |  |
| PC2.temp * group [2] |  |  |  |  |  |  |  |  |  |  | 0.70 | 0.64 |  |  |
| PC2.temp * group [3] |  |  |  |  |  |  |  |  |  |  | 0.36 | 0.59 |  |  |
| PC2.temp * group [4] |  |  |  |  |  |  |  |  |  |  | 0.38 | 0.59 |  |  |
| PC1.precip * group [2] |  |  |  |  |  |  |  |  |  |  |  |  | -0.39 | 0.32 |
| PC1.precip * group [3] |  |  |  |  |  |  |  |  |  |  |  |  | -0.52 | 0.32 |
| PC1.precip * group [4] |  |  |  |  |  |  |  |  |  |  |  |  | -0.36 | 0.33 |
| PC2.precip * group [2] |  |  |  |  |  |  |  |  |  |  |  |  | 0.33 | 0.21 |
| PC2.precip * group [3] |  |  |  |  |  |  |  |  |  |  |  |  | 0.30 | 0.21 |
| PC2.precip * group [4] |  |  |  |  |  |  |  |  |  |  |  |  | 0.42 | 0.22 |
| Model $R^2$ / $R^2$ adjusted | 0.113 / 0.095 | | 0.292 / 0.281 | | 0.305 / 0.280 | | 0.290 / 0.261 | | 0.335 / 0.296 | | 0.332 / 0.285 | | 0.358 / 0.314 | |

Model 5 included all environmental variables, group, and interactions among environmental variables. In this model, two of the groups, *R. integrifolia* × *ovata* and Californian *R. ovata*, showed a significantly positive relationship with leaf area (coef = 0.13, se = 0.04, p < 0.01; coef = 0.22, se = 0.07, p < 0.01, respectively), while the same two interactions were significant as in Model 4. Model 6 included all environmental PCs, group, and all group × temperature PC interactions. Here, PC1_precip_ had a significant negative association with leaf area (coef = -0.08; se = 0.03, p < 0.05), while no other terms were significant. Model 7 was identical to model 6 but instead included all group × precipitation PC interactions; none of the terms were significant, as was the case with Model 8 (the ‘full’ model), which included all interactions among group × environmental PCs. Thus, there was no further explanatory power by including all interaction terms.

### 3.3 Population genetics

Fig. 3 reveals a total of 42 phased nuclear ITS haplotypes and 13 plastid haplotypes (combined *ndhC-trnV* and *rpl16-rps3*). Immediately evident from Fig. 3 is that several ITS haplotypes are shared among individuals containing *R. integrifolia, R. ovata*, and localities with putatively introgressed individuals. A single ITS haplotype is shared by most individuals of *R. integrifolia*, with several closely related haplotypes (Fig. 3; 4). A similar pattern of sharing was observed for plastid DNA (Figs. 3; 4), with greater evidence of structuring among localities than for ITS. Five plastid DNA haplotypes are present among individuals sampled from *R. integrifolia* localities, while ten are present among individuals from *R. ovata* from California. A single plastid haplotype is shared by all AZ individuals of *R. ovata*, while six haplotypes spanning most of the network are shared among individuals from localities containing *R. integrifolia x ovata*.

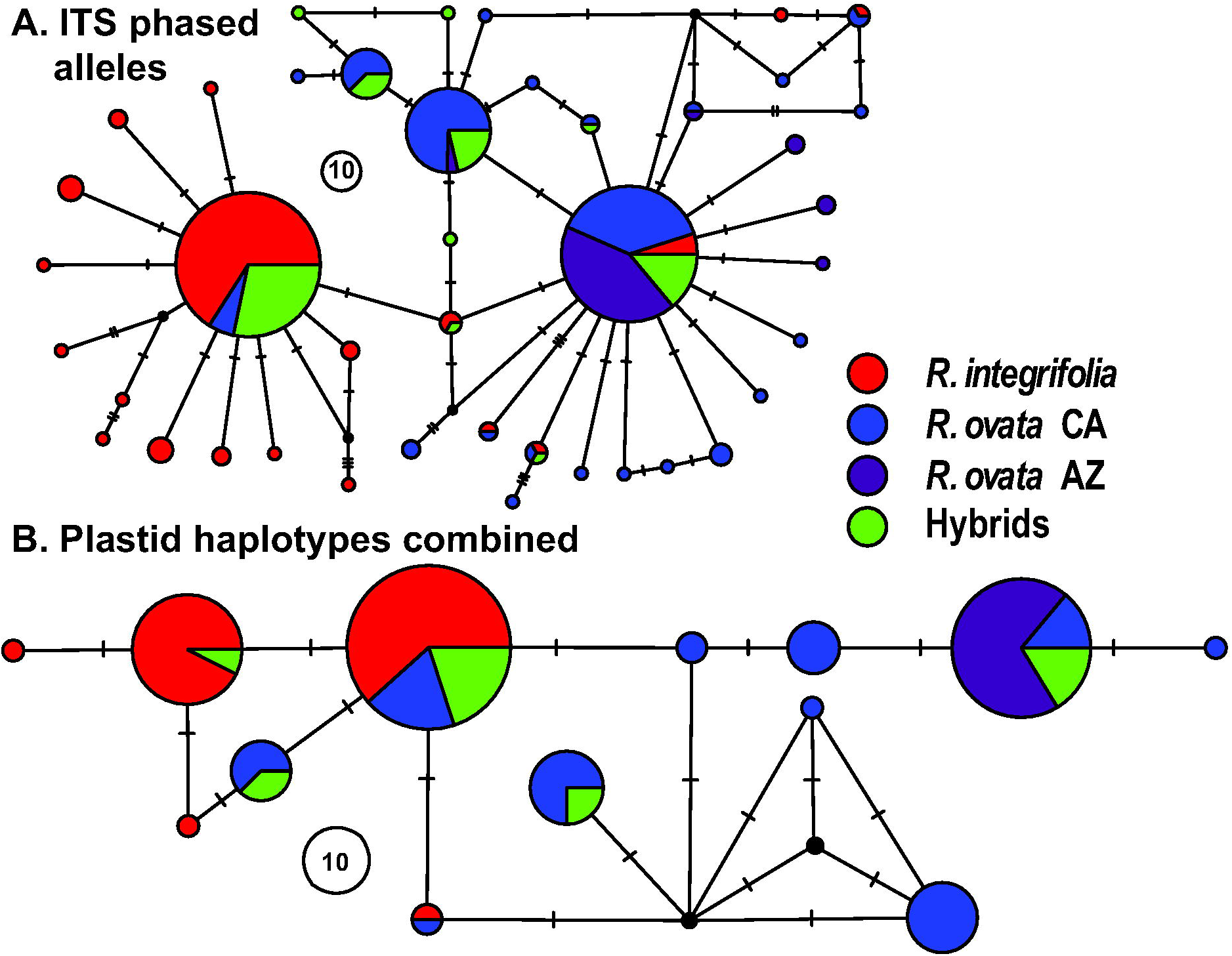

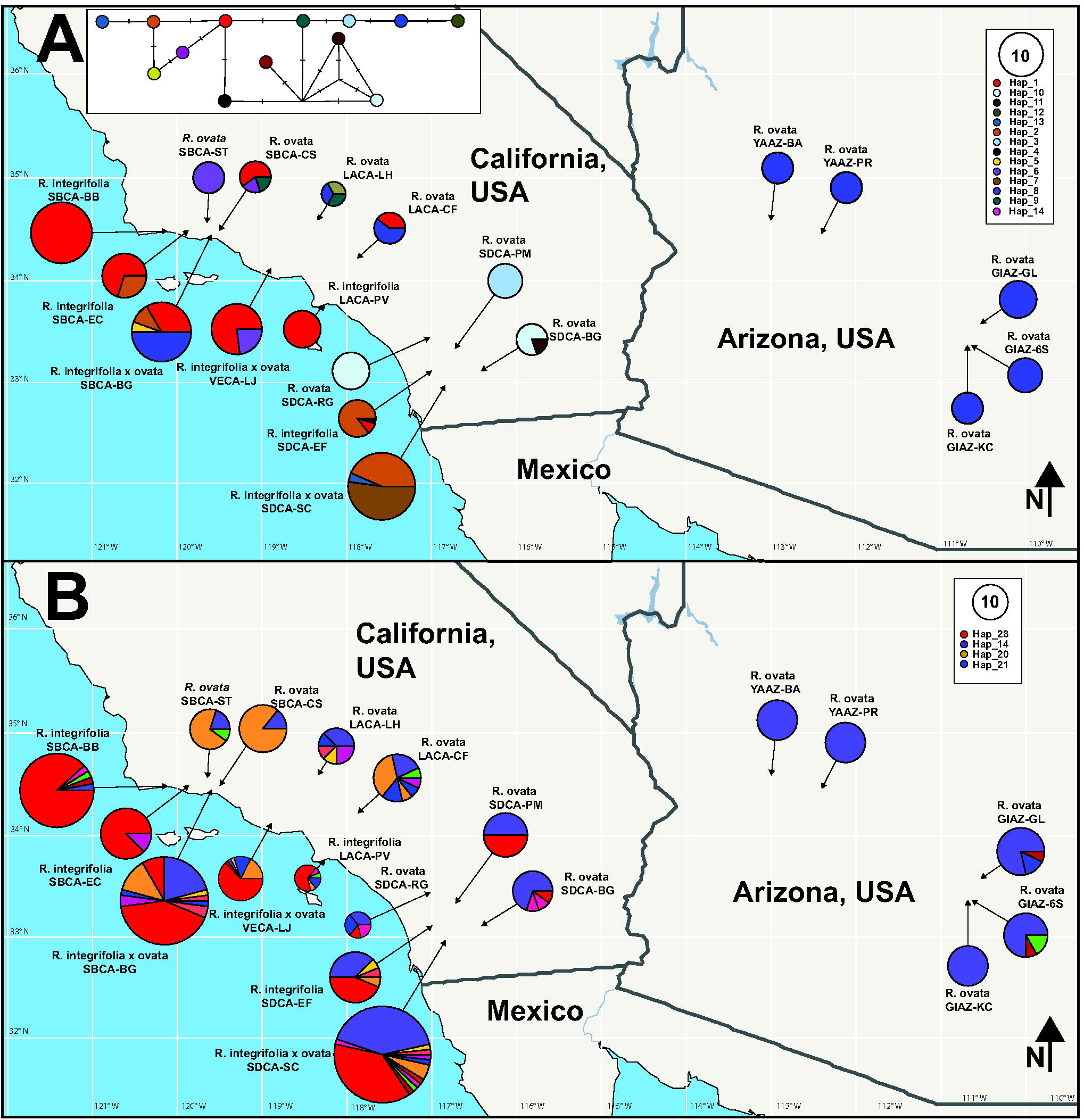

Overall, ITS and plastid haplotype richness is highest for *R. integrifolia × ovata* localities (Table 4), followed by *R. ovata* (CA), *R. integrifolia*, and *R. ovata* (AZ), respectively. Nucleotide diversity is generally the highest in Californian *R. ovata* (π_plastid_ range = 0.000-0.700; π_ITS_ range = 0.234-0.780) and *R. ovata × integrifolia* localities (π_plastid_ range = 0.385-0.660; π_ITS_ range = 0.576-0.771), and lowest in AZ localities of *R. ovata* (π_plastid_ = 0.000; π_ITS_ range = 0.000-0.439).

**Table 4.** Nucleotide and haplotype diversity among sampling localities. π-pt = pairwise nucleotide diversity for plastid DNA; A-pt = the number of plastid haplotypes; n-pt = sample size for plastid DNA; π-ITS = pairwise nucleotide diversity for ITS; A-ITS = the number of ITS haplotypes; n-ITS = sample size for ITS.

| Species present | Locality code | π-pt | A-pt | n-pt | π-ITS | A-ITS | n-ITS |
| --- | --- | --- | --- | --- | --- | --- | --- |
| *R. integrifolia* | LACA-PV | 0.000 | 1 | 7 | 0.593 | 5 | 14 |
| *R. integrifolia* | SBCA-EC | 0.467 | 2 | 10 | 0.233 | 2 | 16 |
| *R. integrifolia* | SDCA-EF | 0.286 | 2 | 7 | 0.700 | 5 | 16 |
| *R. integrifolia* | VECA-BB | 0.000 | 1 | 19 | 0.225 | 4 | 34 |
| *R. integrifolia* × *ovata* | SBCA-BG | 0.660 | 4 | 18 | 0.771 | 10 | 48 |
| *R. integrifolia* × *ovata* | VECA-LJ | 0.385 | 2 | 13 | 0.576 | 6 | 40 |
| *R. integrifolia* × *ovata* | SDCA-SC | 0.561 | 3 | 23 | 0.691 | 12 | 58 |
| *R. ovata* | SBCA-CS | 0.700 | 3 | 5 | 0.234 | 2 | 14 |
| *R. ovata* | SBCA-ST | 0.650 | 1 | 5 | 0.511 | 3 | 10 |
| *R. ovata* | SDCA-BG | 0.800 | 2 | 5 | 0.533 | 4 | 10 |
| *R. ovata* | LACA-CF | 0.500 | 2 | 5 | 0.846 | 7 | 10 |
| *R. ovata* | LACA-LH | 1.000 | 3 | 3 | 0.857 | 5 | 8 |
| *R. ovata* | YAAZ-PR | 0.000 | 1 | 5 | 0.000 | 1 | 10 |
| *R. ovata* | YAAZ-BA | 0.000 | 1 | 5 | 0.000 | 1 | 10 |
| *R. ovata* | GIAZ-GL | 0.000 | 1 | 7 | 0.384 | 3 | 14 |
| *R. ovata* | GIAZ-6S | 0.000 | 1 | 6 | 0.439 | 3 | 12 |
| *R. ovata* | GIAZ-KC | 0.000 | 1 | 5 | 0.000 | 1 | 10 |
| *R. ovata* | SDCA-PM | 0.000 | 1 | 5 | 0.545 | 2 | 12 |
| *R. ovata* | SDCA-RG | 0.000 | 1 | 7 | 0.78 | 4 | 14 |

AMOVA revealed significant structure among the four groupings for ITS (Table 5; Φ_CT_ = 0.369, % variation = 55.837, p < 0.001) and plastid DNA (Φ_CT_ = 0.364, % variation = 31.644, p < 0.001). Both ITS and plastid DNA showed significant structure among localities within groupings, with plastid DNA having higher levels of structuring among localities than nuclear ITS (Table 4; for plastid DNA Φ_SC_ = 0.557, % variation = 57.023, p < 0.001; for ITS Φ_SC_ = 0.123, % variation = 7.251, p < 0.001). This pattern is further illustrated by pairwise Φ_ST_ comparisons among localities (Fig. 5). Pairwise Φ_ST_ values are generally significant and relatively high when comparing *R. integrifolia* with *R. ovata* from CA and AZ localities, but lower compared to *R. integrifolia × ovata* localities. A similar overall pattern is observed for ITS, but in general pairwise Φ_ST_ values are lower, reflecting the results from AMOVA of weaker overall structure among localities relative to plastid DNA. N_ST_ and G_ST_ values were 0.786 and 0.696 for plastid DNA, and 0.431 and 0.377 for ITS, respectively. For both plastid DNA and ITS the differences between N_ST_ and G_ST_ were significant (0.090, p < 0.001; 0.054, p < 0.001, respectively).

**Table 5.** Analysis of Molecular Variance (AMOVA, in Arlequin). ‘df’ = degrees of freedom, ‘SS’ = sum of squares, ‘SD’ = standard deviation, ‘% variation’ = percent of total variation explained, ‘Φ**’** refers to the analog of inbreeding coefficients (F-statistics) for haploid data, and ‘p’ = significance of differentiation at each hierarchical level.

| **Plastid** | **df** | **SS** | **SD** | **%**<br>**variation** | **$\Phi$** | **p** |
| --- | --- | --- | --- | --- | --- | --- |
| **Within localities ($\Phi_{ST}$)** | 2 | 120.701 | 0.473 | 11.332 | 0.719 | < 0.001 |
| **Among localities within groupings ($\Phi_{SC}$)** | 16 | 331.470 | 2.382 | 57.023 | 0.557 | < 0.001 |
| **Among groupings ($\Phi_{CT}$)** | 152 | 200.881 | 1.322 | 31.644 | 0.364 | < 0.001 |
| **Total** | 170 | 653.053 | 4.176 | n/a |  | n/a |
| **ITS** |  |  |  |  |  |  |
| **Within localities ($\Phi_{ST}$)** | 2 | 284.812 | 1.142 | 36.875 | 0.391 | < 0.001 |
| **Among localities within groupings ($\Phi_{SC}$)** | 16 | 90.333 | 0.225 | 7.251 | 0.123 | < 0.001 |
| **Among groupings ($\Phi_{CT}$)** | 345 | 596.835 | 1.730 | 55.873 | 0.305 | < 0.001 |
| **Total** | 363 | 972.000 | 3.0 | n/a |  | n/a |

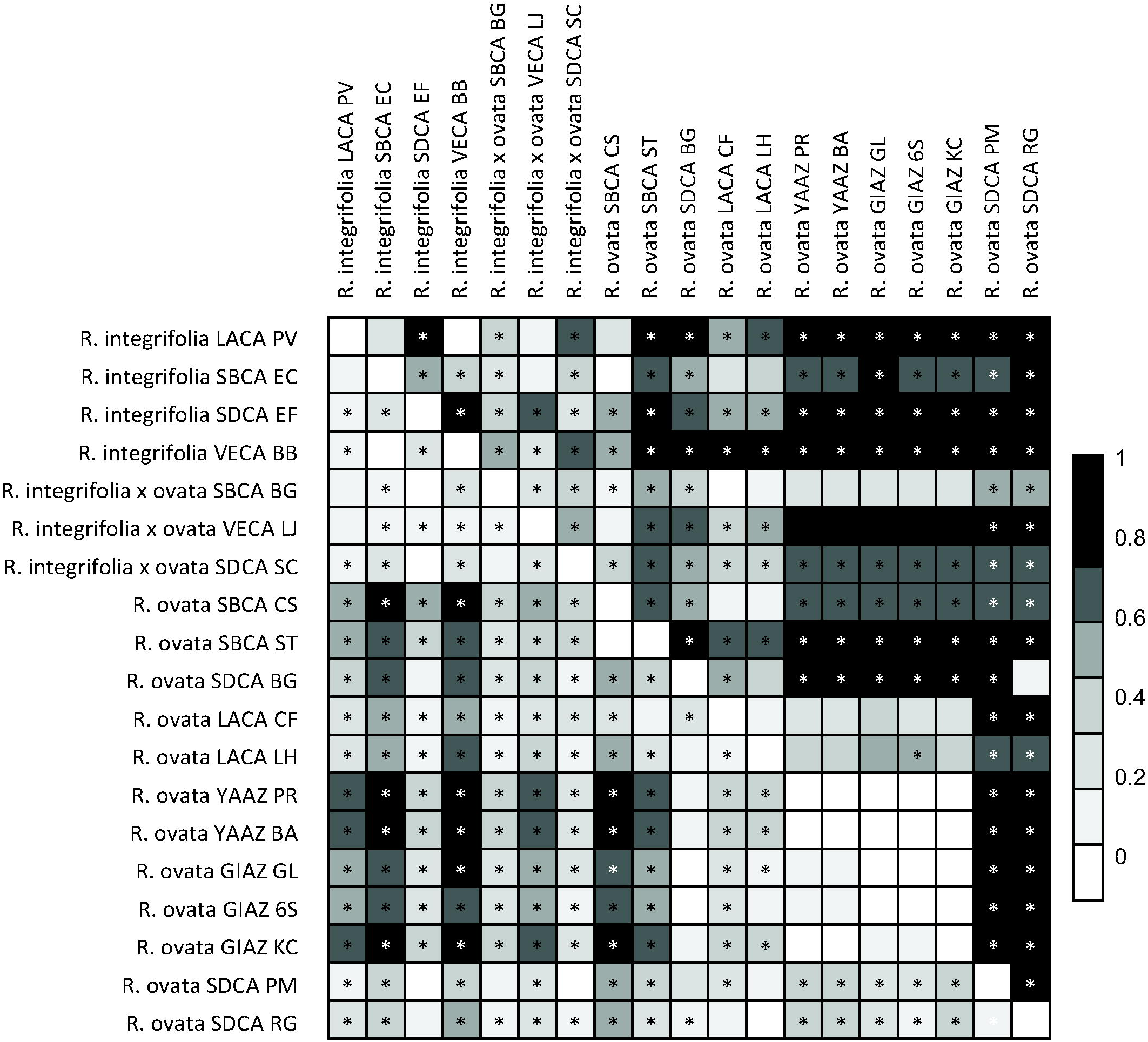

P_ST_-Φ_ST_ comparisons reveal differentiation in leaf morphology among localities, using the first two Principal Components as composite proxies (Fig. 6). P_ST_ was estimated for a range of values of c/h^2^, which corresponds to the ratio of additive genetic variation in morphology among localities (c) to that among individuals (h^2^, i.e. narrow-sense heritability). Using the estimated global Φ_ST_ values of 0.441 for ITS and 0.684 for plastid DNA, the thresholds for c/h^2^ that correspond to P_ST_ > Φ_ST_ are approximately 0.1/0.4 for PC1, and 0.2/0.5 for PC2 (for plastid DNA and ITS, respectively).

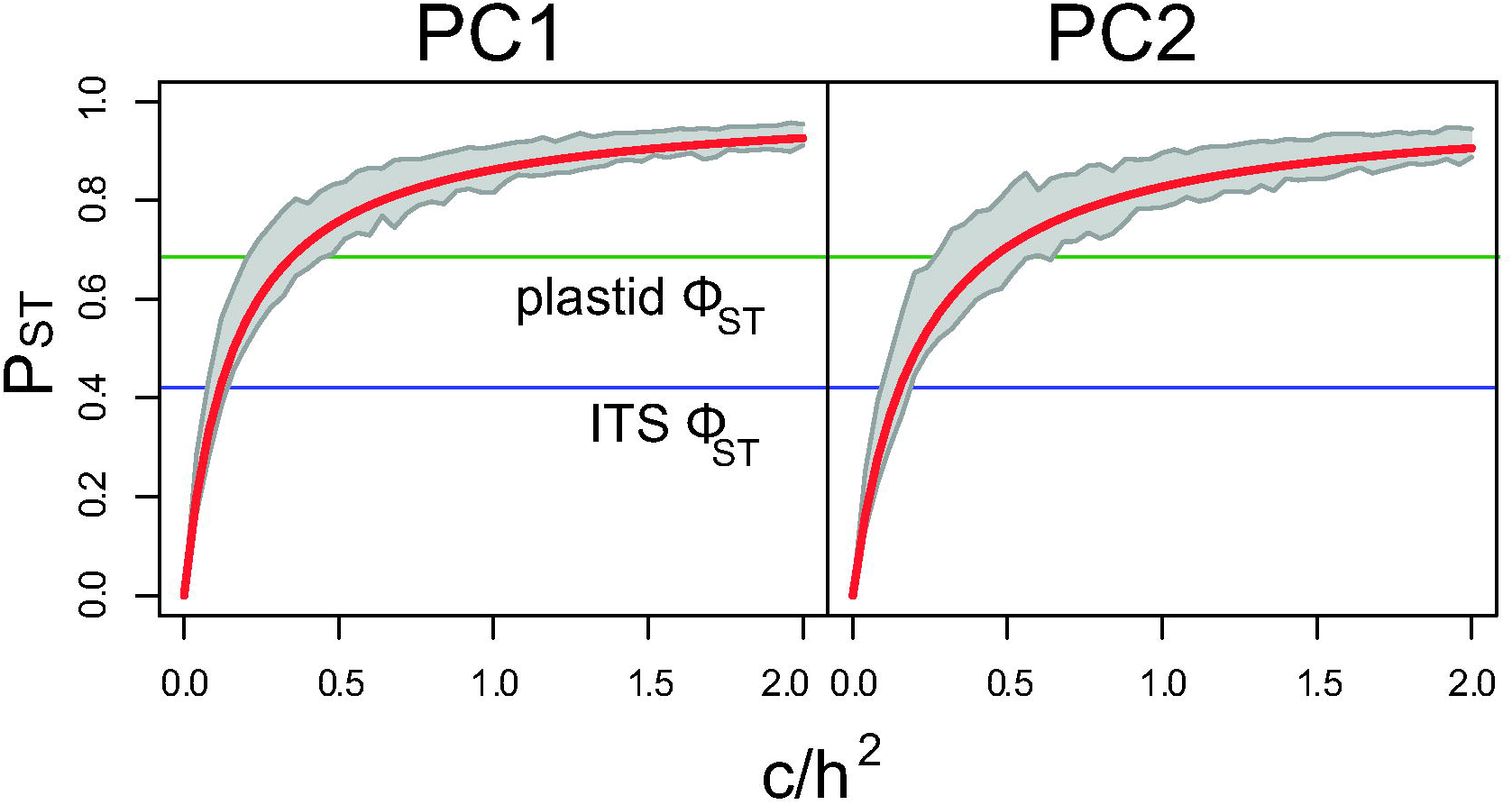

### 3.4 Species distribution modeling

Filtering of occurrence data with CoordinateCleaner retained 503 of 1,209 occurrence points for *R. integrifolia* and 852 of 1,523 occurrence points for *R. ovata*. PCA of all 19 environmental variables revealed that the localities sampled only represent a subset of the total multivariate environmental niche space represented by a larger collection of herbarium records from GBIF (Fig. S6). PC1 explains 46.3% of the total variation in environmental variables, and is positively associated with mean diurnal range, temperature seasonality, and temperature/precipitation in the driest/warmest quarters; PC1 is negatively associated with precipitation seasonality, isothermality, and temperature during the coldest month/quarter (Table S7). PC2 is positively associated precipitation (annual, wettest month/quarter, and coldest quarter), and negatively associated with temperature during the warmest/wettest quarters. Sampling localities for *R. integrifolia × ovata* fall at intermediate positions between those for *R. integrifolia* and *R. ovata* from California, while Arizonan *R. ovata* occupy a more distinct portion of multivariate space associated with warmer, more seasonal habitats (Fig. S6).

Pearson analysis detected autocorrelation between many bioclimatic predictor variables. The following variables passed this autocorrelation filter and were used to infer SDMs: annual mean temperature (1), mean diurnal range (2), isothermality (3), mean temperature of warmest quarter (8), mean temperature of driest quarter (9), annual precipitation (12), precipitation of driest month (14), precipitation seasonality (15), precipitation of warmest quarter (18), and precipitation of coldest quarter (19). MaxEnt inference of species distribution models for both *R. integrifolia* and *ovata* accurately recapitulated their present ranges and showed that the historical range of *R. ovata* has been highly influenced by climate change over the past ∼22,000 years (Fig. 7). The average training AUC for the ten replicate MaxEnt runs using contemporary climatic variables was 0.988 and 0.977 for *R. integrifolia* and *R. ovata*, respectively. Standard deviation for training AUC was less than 0.001 for each replicate.

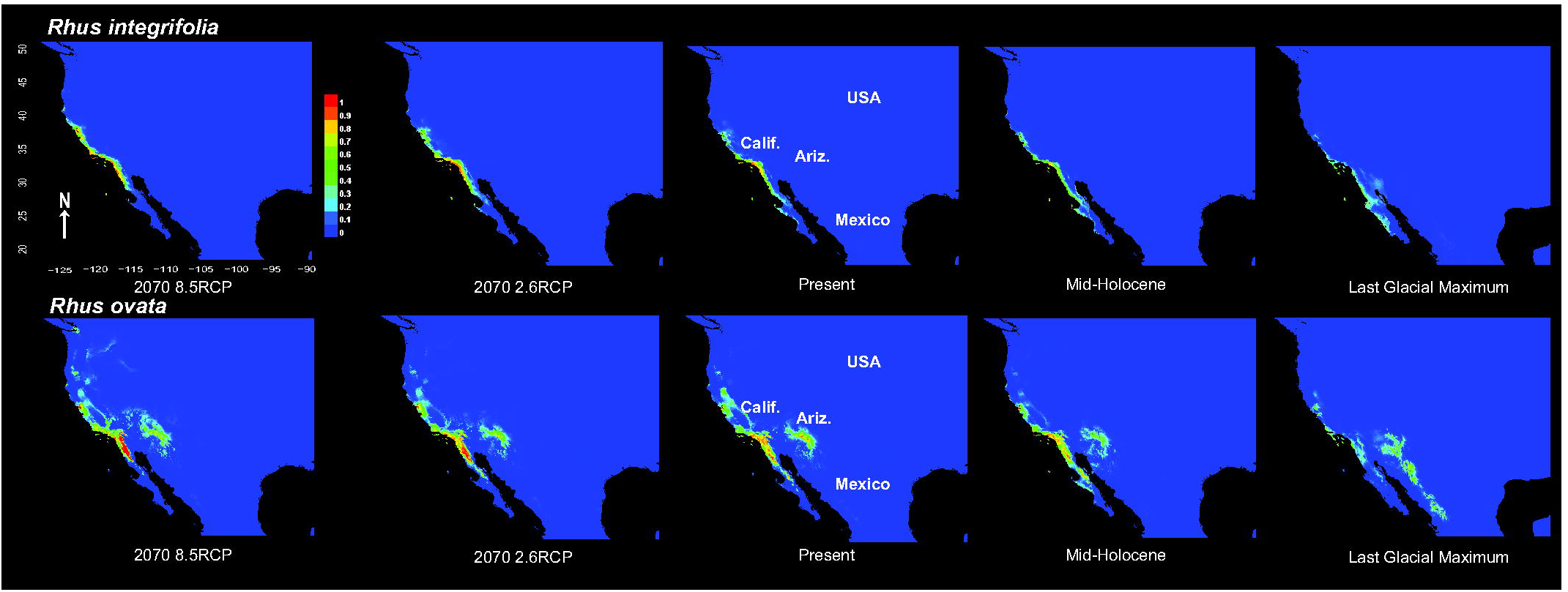

During the LGM, areas projected to be most habitable for *R. integrifolia* were nearly restricted to Baja California (Fig. 7). Contrastingly, most of the habitable range of *R. ovata* during the Last Glacial Maximum was inferred to exist in a relatively narrow band extending from present day Arizona south to Durango, Mexico. However, the reconstructed SDM for *R. ovata* under a Mid- Holocene climate reflects a pronounced range shift, with most of the habitable area overlapping that of *R. integrifolia* in Baja California. *Rhus integrifolia* was inferred to gradually shift northward over time, from Baja California to the central Californian coast. *Rhus ovata* populations were inferred to have a decreasing probability of occurrence over time in Arizona, with increasing probability in coastal habitats. Perhaps most interestingly, the area including present day Arizonan populations of *R. ovata* was inferred to be habitable since the LGM, suggesting that these populations may have become isolated from the remainder of the range sometime between 22 and 8 kya. RTR-MO analyses failed to reject niche equivalency of present ranges (p = 0.49). Our SDM projections suggest that the overlap in habitable range for these two species is likely to increase through 2070, regardless of RCP scenario, as the range of *R. integrifolia* tracks northward along the California coast, and the projected habitability of interior regions declines for *R. ovata*.

## 4 DISCUSSION

Here we present a broad geographic analysis of two ecologically important, hybridizing species, *Rhus integrifolia* and *R. ovata*. We present morphological and molecular evidence for distinctness among the two species, but with clear intermediacy in morphology and haplotype sharing for several localities where they are sympatric. We found population structure in morphology that outranks neutral genetic structure, which may be driven by either environmental plasticity or local adaptation. There is weak evidence for a negative association between overall precipitation and leaf area, but this association is amplified with increasing overall temperature, and offset by increasing temperature variation. Lastly, we infer that range shifts over the last ∼20,000 years likely reflect periods of both sympatry and allopatry, and that future climate conditions will favor increased range overlap among the two species in coastal regions and a decreased probability of *R. ovata* occurrence in Arizona.

### 4.1 Morphology

There is a clear difference among *R. integrifolia* and *R. ovata* based on extensive sampling of leaf morphological characters, and localities containing both parental species display intermediate features based on multivariate analyses of morphology (Fig. 2). This intermediacy is likely to be indicative of the degree to which the genome of each individual is introgressed, i.e. how much of the genome is represented by each parental species. Based on our sampling, introgressant populations occur at mid-elevation localities, between approximately 300-400 m above sea level (Fig. S1). The observed patterns of morphological intermediacy reflect those in other systems including hybrid introgression in California (e.g. Dorado et al., 1991; Albert et al., 1997; Dodd and Afzal-Rafii, 2004).

It is unclear what, if any, positive or negative fitness consequences there may be for introgressive hybridization in this species complex. It remains to be tested whether selection against alleles from the other species may limit the spread of these alleles between parental species in the allopatric portion of their ranges, or if intermediate hybrid populations serve as a bridge for the exchange of adaptive variation among the two species (Barton and Hewitt, 1985; Rieseberg and Burke, 2001). While we found morphological differentiation (P_ST_) to outrank neutral genetic differentiation (Φ_ST_) above a certain heritability threshold (i.e. c^2^/h = 0.2 and 0.4 for ITS and plastid DNA, respectively; Fig. 6), it remains unclear whether this is caused by extensive phenotypic plasticity or adaptive genetic variation.

### 4.2 Leaf area and environmental variation

Leaf area has long been recognized as an important functional trait at the inter- and intraspecific levels (e.g. Osnas et al., 2013; 2018). Young (1974) noted a possible relationship between latitude and leaf size in *R. ovata*, based on limited sampling of three localities. We have explicitly tested this pattern in the context of abiotic environmental variation, which represents a more accurate approach than using latitude, longitude, and elevation as proxies for environmental variation. Our analysis of 19 sampling localities highlights differences among groupings (e.g. *R. ovata, R. integrifolia*, and introgressant populations) as the primary determinants of leaf area (as corroborated by Fig. 2), but also suggests an association between overall precipitation (PC1_precip_) and leaf area (Table 3). Although weak, this association is negative, with smaller leaves at localities with higher overall precipitation. However, there is interaction between overall temperature + overall precipitation, suggesting these two factors together may at least partially influence leaf area. This association is offset by temperature variation; i.e., warmer, wetter areas tend to harbor populations with smaller leaves, but local climates with more drastic temperature swings show an opposite trend (Table 3).

Given that that species in xeric regions tend to have leaves that are small (e.g. Givnish et al., 1979), succulent, dissected, or even absent (e.g. cacti, euphorbias), it seems counterintuitive that leaf area would decrease with increasing precipitation in this species complex. In fact, we found the opposite of what has been observed across many other species, as larger leaves are thought to shed heat more slowly due to a larger boundary layer in hot, arid environments, thus posing a risk for overheating and extensive evaporative water loss (Schuepp, 1993). *Rhus ovata* has thick, waxy leaves that fold adaxially along the midrib during the hottest, driest months, which likely represents an adaptation to seasonally extreme heat and aridity in chaparral habitats (e.g. Herbert and Larsen, 1985). However, *R. ovata* experiences temperatures below freezing, especially at high-elevation inland localities (Boorse et al., 1998; Montalvo, 2017), and thus the relationship observed between leaf area, precipitation, and temperature may reflect a complex tradeoff between climatic extremes. Other factors not included in our analysis, such as soil fertility, grazing pressure, and density-dependence may interact with abiotic climatic factors, and provide a clearer picture of the determinants of leaf area in *Rhus*. There are some localities at which *R. integrifolia, R. ovata*, and their hybrids occur within close proximity, with steep elevational gradients (e.g. in the Santa Monica Mountains of California). These areas provide the ideal grounds for ‘natural experiments’ to study the dynamics of leaf area as it relates to putative adaptations to temperature, precipitation, and hybridization.

### 4.3 Population genetics

Nuclear ITS and two plastid markers display clear patterns of allele sharing at localities with hypothesized introgression, reflecting a congruent pattern to that of morphological intermediacy (Figs. 2-4). However, these shared haplotypes are not restricted to localities in which plants display intermediate morphologies; indeed, the two most common ITS and plastid haplotypes are widespread, being shared at localities corresponding to either morphologically distinct *R. integrifolia* or Californian *R. ovata*. Within *R. ovata*, which is disjunct from California to Arizona, four of six Californian localities contain multiple plastid haplotypes, whereas all Arizonan populations contain a single, identical haplotype. Three of five Arizonan localities contain a single, common ITS haplotype. This finding suggests a potential bottleneck, or that smaller effective population sizes in Arizona may be prone to the effects of genetic drift, resulting in overall lower genetic diversity there. Even so, ITS reveals that some haplotypes are shared widely across the network, differing somewhat from the pattern based on plastid DNA. For example, the most common ITS haplotype among *R. integrifolia* localities is also found in CA and AZ localities of *R. ovata*. Likewise, the most common ITS haplotype in AZ localities is also found in Californian *R. ovata* localities, and even in *R. integrifolia*.

Overall, weaker population structure for ITS than for plastid DNA could be driven by two factors: larger effective population sizes and unsorted ancestral polymorphism for nuclear DNA than for organellar DNA (Palumbi and Baker, 1994; McCauley, 1995; Avise, 2000; Hare, 2001; Palumbi et al., 2001), or greater interpopulation dispersal range for pollen vs. seeds (e.g. Ennos, 1994; Hamilton, 1999; Kartzinel et al., 2013). Both *Rhus* species are predominantly pollinated by bees (Young, 1972; Moldenke and Neff, 1974), while seeds are dispersed by mammals and birds (Lloret and Zedler, 1991; Rowe and Blazich, 2008). Our findings are congruent with a hypothesis in which barriers to pollen dispersal are low among populations of both species and between them, albeit with decreased fecundity for ‘hybrid’ individuals, as observed in previous experimental crosses (Young, 1972). Negative fecundity barriers may be overcome if pollen flow occurs frequently enough over long enough distances. Seed dispersal, on the other hand, may be limited by successful recruitment (e.g. Dunne and Parker, 1999; Arrieta and Suarez, 2006), which may be amplified if avian or mammalian vectors travel long distances dispersing seeds at unfavorable localities with different local environmental conditions. This is especially pertinent if local conditions (soil, temperature, moisture, fire regime, frost formation) vary enough across sites such that environmental differences between source and sink sites suppress successful colonization by immigrant propagules, which may not be able to compete in new localities. Boorse et al. (1998) conducted the only study documenting putative evidence for local adaptation in *R. ovata*: plants from a site with lower minimum winter temperatures were significantly less susceptible to freezing damage than plants from a warmer site in the same region of coastal California. Furthermore, leaves of seedlings were much more susceptible to freezing than those from adult plants, possibly representing a barrier to successful establishment by propagules from nearby warmer habitats. Reciprocal transplants using seeds of *R. integrifolia, R. ovata*, and their hybrids could yield useful data on whether recruitment is limited by local adaptation, and whether this has implications for population structure as it relates to seed dispersal and successful establishment.

Unsorted ancestral polymorphism would seem unlikely in this case to be the sole explanation for widely shared ITS haplotypes, given the previously estimated divergence time of ∼3.1 mya between *R. integrifolia* and *R. ovata* (Yi et al., 2004). Widespread pollen flow (both historical and contemporary), potential environmental barriers to external propagule recruitment in established populations, and differences in plastid vs. nuclear DNA effective population sizes may all contribute to the discrepancy in population structure among localities. Genome-scale data would allow several hypotheses to be tested regarding demographic history, patterns of gene flow between the two species and their introgressants, and the adaptive value (if any) of introgression. For example, a comparison of which regions of the genome have experienced higher or lower rates of introgression would be particularly insightful, especially in the context of putatively adaptive or maladaptive introgression (e.g. Grant and Grant, 1998; Rieseberg and Burke, 2001). Furthermore, these data would provide the level of resolution needed for the construction of historical demographic models under ancestral and contemporary gene flow vs. retained ancestral polymorphism.

### 4.4. Species distribution modeling

Our species distribution models accurately capture the ranges of both species despite the lack of significant niche differentiation between them. The lack of niche differences between our MAXENT distribution models based on the RTR-MO test likely reflects the existence of hybrid/introgressed populations of *R. integrifolia* and *R. ovata* and their high degree of niche similarity in coastal regions (Fig. 7). Hybrid populations present a problem for species distribution models; specimens from GBIF are identified either as *R. integrifolia* or *R. ovata* but do not include information on hybrid status. A meticulous analysis of morphology, and perhaps even genetic variation from herbarium specimens might improve resolution of future species distribution models in the *R. integrifolia*-*ovata* complex, by allowing populations with evidence of hybrid introgressants to be treated as a separate category. However, even this approach may be an oversimplification, because the degree to which each population is introgressed is likely to vary across regions of overlap among the parental species (Arnold, 1997), and thus the application of a hybrid index on a continuous scale may be more appropriate (e.g. Cullingham et al., 2012).

Based on our models, we infer a northward shift in the distribution of *R. integrifolia*, as well as a higher concentration of occurrence immediately along Californian coast forecasted for 2070, for both the best- and worst-case projections of future atmospheric CO_2_ levels (Fig. 7). Thus, the distribution of *R. integrifolia* is likely to be forced by climate change to shift into one of the most densely populated regions in North America, where human development continues to compromise coastal habitats. For *R. ovata* we infer a similar northward shift for interior populations in Arizona, and a gradually decreasing occurrence probability in that region. Our models also infer an increased occurrence probability along the Californian coast and in northern Baja California for *R. ovata*. Riordan et al. (2018) used species distribution modeling in several southern Californian chaparral plant species and predicted that suitable habitat would generally remain stable in the future for *R. ovata*. However, they also predicted that habitat gains in low-elevation areas for *R. ovata* will likely coincide with future human development, and that population fragmentation is predicted to increase, with implications for gene flow and local adaptation.

Taken together, our 2070 forecast indicates that both species will be forced into a higher degree of sympatry, possibly increasing the occurrence of introgression. A worst-case scenario would include decreasing occurrence probability in the allopatric part of the range for *R. ovata* (i.e. Arizona), coupled with the prediction that *R. integrifolia* and *R. ovata* will be pushed into coastal regions. Though it is impossible to predict exactly what will happen in the future, global climate change may contribute to the erosion of locally adapted variants and possibly species boundaries by increasing the frequency of hybridization among these two species. This finding highlights the need for investigation of the importance of local adaptation within each species, and the adaptive consequences of potentially increased levels of hybridization among them, e.g. via common garden experiments, reciprocal transplants with experimental crosses, and genomic analysis.

### 4.5 Taxonomic implications

Hybridization has long presented challenges for species delimitation, especially as it relates to the Biological Species Concept (e.g. Mallet, 2005). While *R. ovata* and *R. integrifolia* are clearly distinct in regions of allopatry, the same cannot be said within regions of sympatry. The two species become nearly indistinguishable in the latter, forming a continuous gradation in morphology, genetic variation, and niche overlap. The evidence presented here does not warrant specific taxonomic changes, but it does suggest that hybrid status should be considered when depositing new collections in herbaria or other specimen databases. Furthermore, popular instruments for the ‘crowdsourcing’ of species occurrence data could be used to help distinguish among more “pure” forms of each species and their hybrid introgressants, at least for contemporary observation records. For example, using the application iNaturalist (https://www.inaturalist.org/), there are 170 records of *R. integrifolia × ovata* (most likely an underestimate of their actual abundance), 5,715 of *R. integrifolia*, and 3,591 of *R. ovata* (last accessed April 14, 2020). While these types of observations have obvious biases (i.e. they tend to be clustered near population centers or in public lands; e.g. Dickinson et al., 2012), if properly verified either visually by an expert or via machine learning (e.g. Priya et al., 2012; Wilf et al., 2016; Kaur and Kaur, 2019) they may ultimately improve distribution models for hybridizing species, including *R. integrifolia* and *R. ovata*.

The acquisition of genome scale variation is now feasible for many researchers and should prove to be particularly informative in determining patterns of gene flow among these two species (Taylor and Larsen, 2019). Such data will allow the detection of adaptive variants, and regions of the genome that experience higher or lower levels of gene flow than regions under neutral expectations (Whitney et al., 2010). Ultimately this type of genomic information could be used to infer the relative importance of selection in maintaining species boundaries in the face of frequent gene flow (Rieseberg and Burke, 2001; Feder et al., 2012; Suarez Gonzalez et al., 2018).

## 5 CONCLUSIONS

We investigated morphological, genetic, and environmental variation in two ecologically important, hybridizing species of *Rhus* in the southwestern USA. Our findings revealed morpho-genetic distinctness among parental species but intermediacy at localities where hybridization is hypothesized to occur. We further found morphological differences among plants from different localities that outrank genetic differentiation, suggesting local adaptation or widespread phenotypic plasticity, and a weak negative relationship between leaf area and overall precipitation. Species distribution models predicted range shifts northward and into coastal habitats for both species, possibly with implications for increased future levels of hybridization. Our study highlights the importance of sampling broadly and integrating morphological, genetic, and ecological niche data, further underscoring the challenges associated with species distribution modeling of hybridizing species. Additional studies using reciprocal transplants of both parental species and their hybrid introgressants, along with genome-wide surveys of variation will help elucidate the relative impacts of gene flow and selection on the maintenance of species boundaries.

## ACKNOWLEDGEMENTS

We thank the USDA Forest Service for permission to collect samples. We also thank Ryan Percifield, Ashley Henderson, and Apoorva Ravishankar (WVU Genomics Core Facility) for support provided to help make this publication possible, and CTSI Grant #U54 GM104942 which in turn provides financial support to the Genomics Core Facility. Funding was provided by the WVU Department of Biology, the WVU Eberly College of Arts and Sciences, and a WVU-Program to Stimulate Competitive Research (PSCoR) award to CB.

