## Supplementary data 1-7 for "Genetic, morphological, and niche variation in the widely hybridizing *Rhus integrifolia-Rhus ovata* species complex": S1 groups x elevation.docx

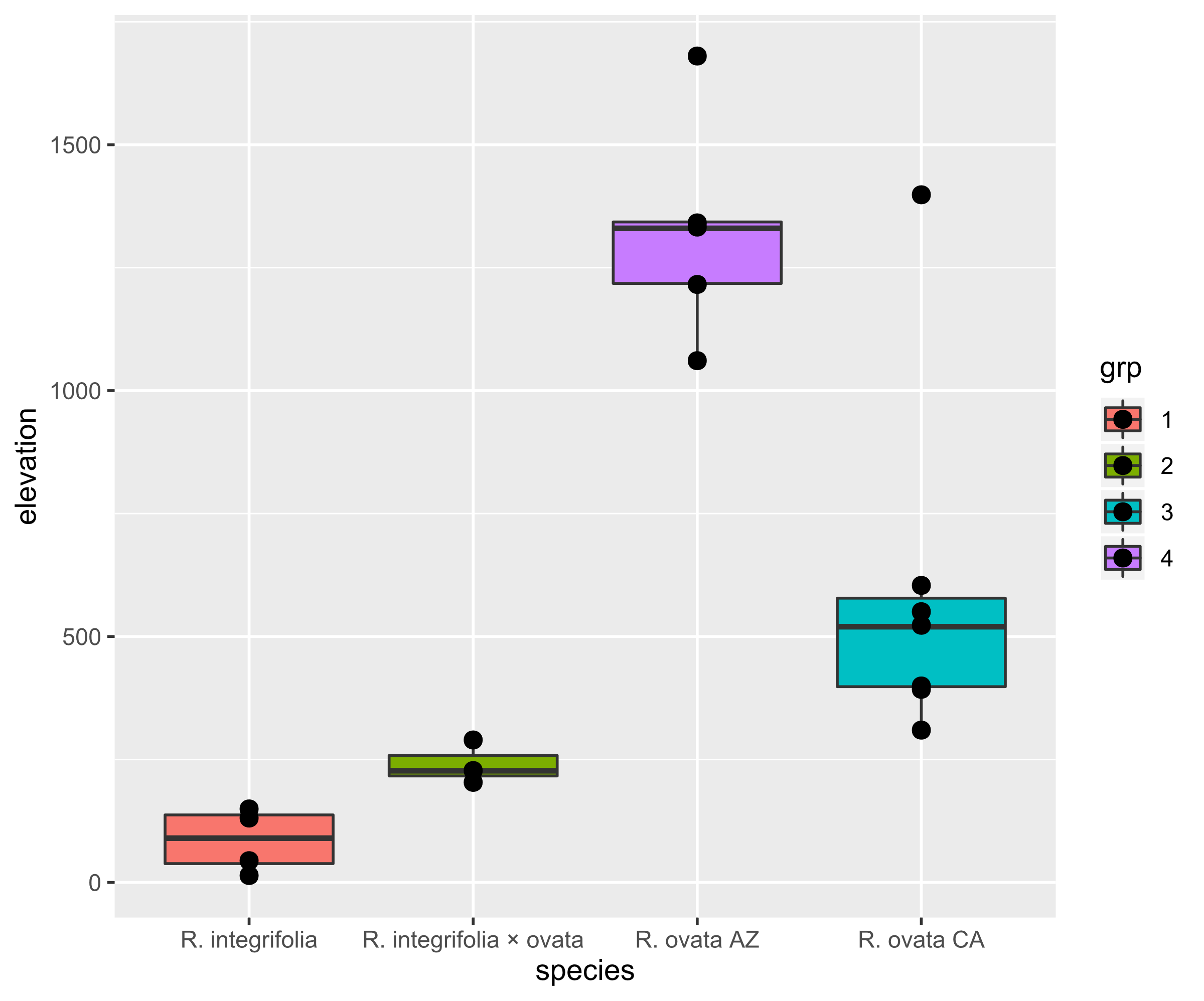

**Fig. S1**. Boxplot of elevation (meters) for sampling localities of: *R. integrifolia* (red), *R. integrifolia × ovata* (green), Californian *R. ovata* (blue), and Arizonan *R. ovata* (purple).
