## Supplementary data 1-7 for "Genetic, morphological, and niche variation in the widely hybridizing *Rhus integrifolia-Rhus ovata* species complex": S2 morph PCA percent variation.docx

**Table S2**. Eigenvalues and % variance explained for principal components analysis of 14 morphological features based on a correlation matrix.

| PC | Eigenvalue | % variance |
| --- | --- | --- |
| 1 | 6.85884 | 48.992 |
| 2 | 1.48807 | 10.629 |
| 3 | 1.10914 | 7.9224 |
| 4 | 0.99657 | 7.1184 |
| 5 | 0.847159 | 6.0511 |
| 6 | 0.667543 | 4.7682 |
| 7 | 0.542979 | 3.8784 |
| 8 | 0.441585 | 3.1542 |
| 9 | 0.362536 | 2.5895 |
| 10 | 0.246141 | 1.7581 |
| 11 | 0.220993 | 1.5785 |
| 12 | 0.125817 | 0.89869 |
| 13 | 0.060439 | 0.43171 |
| 14 | 0.032193 | 0.22995 |
