## Supplementary data 1-7 for "Genetic, morphological, and niche variation in the widely hybridizing *Rhus integrifolia-Rhus ovata* species complex": S3 morphology PCA loadings PC1_2.docx

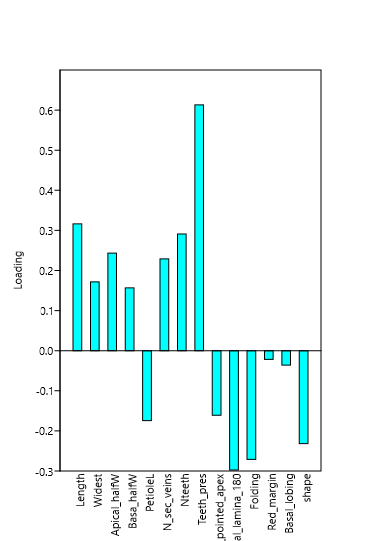

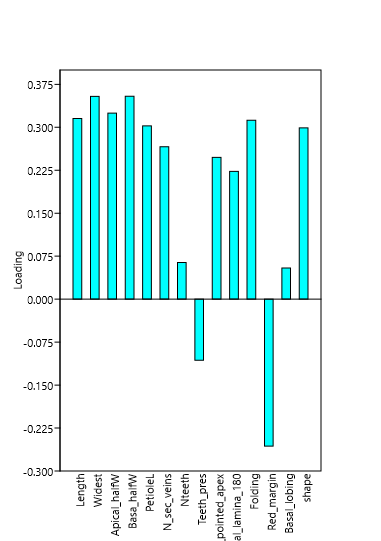

**Fig. S3**. PCA loading scores for 14 morphological features based on principal components analysis using a correlation matrix for PC1 (left) and PC2 (right).
