## Supplementary data 1-7 for "Genetic, morphological, and niche variation in the widely hybridizing *Rhus integrifolia-Rhus ovata* species complex": S4 NEW morph 2 way NPMANOVA.docx

| **Source** | **Sum of squares** | **d.f.** | **Mean square** | **F** | **p** |
| --- | --- | --- | --- | --- | --- |
| **Group (n = 4)** | 3.9858 | 3 | 1.3286 | 9.897 | 0.0001 |
| **Locality (n = 19)** | 1.8853 | 4 | 0.47133 | 3.5111 | 0.0001 |
| **Interaction** | -19.282 | 12 | -1.6069 | -11.97 | 1 |
| **Residual** | 24.432 | 182 | 0.13424 |  |  |
| **Total** | 11.02 | 201 |  |  |  |

**Table S4**. Two-way non-parametric multivariate analysis of variance (NP-MANOVA) based on Gower-transformed distance for mixed data types and 9,999 permutations. “Groups” correspond to: *R. integrifolia*, *R. integrifolia × ovata*, Californian *R. ovata*, and Arizonan *R. ovata*.
