## Supplementary data 1-7 for "Genetic, morphological, and niche variation in the widely hybridizing *Rhus integrifolia-Rhus ovata* species complex": S5 bioclim scree plots and loadings.docx

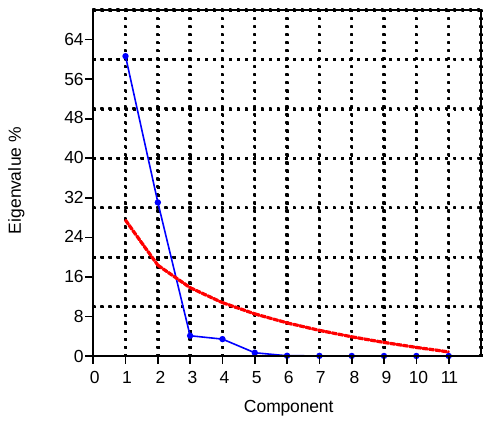

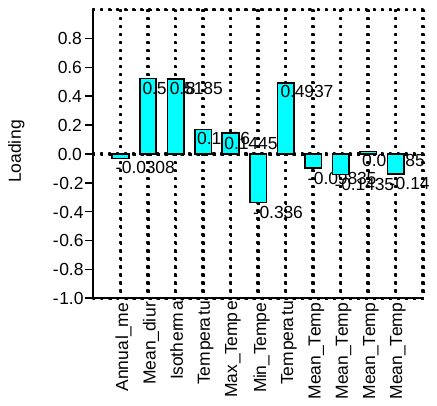

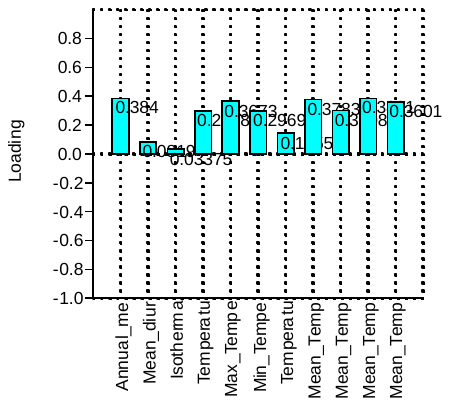

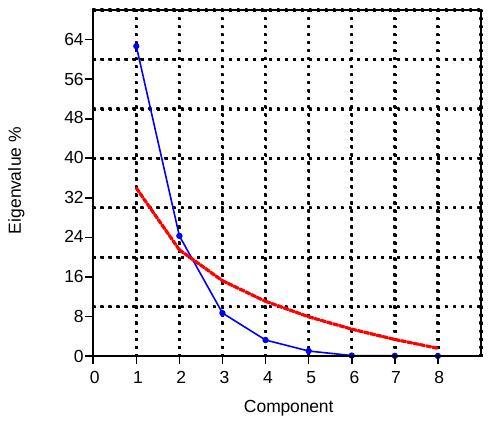

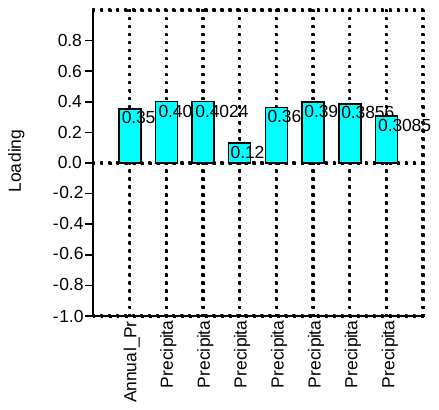

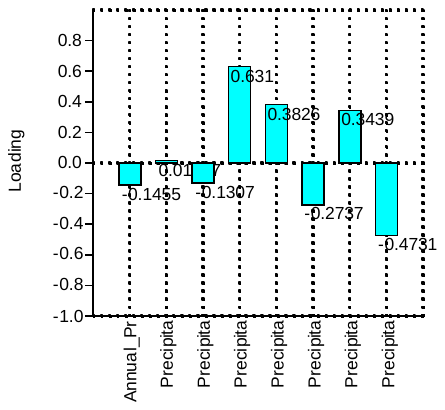

Temperature (BIOCLIM 1-11)

Precipitation (BIOCLIM 12-19)

A

B

C

D

E

F

**Fig. S5**. Scree plot of PCA eigenvalues (blue) and “broken stick” analysis (red) of environmental data based on 19 BIOCLIM variables. Above: Analysis of BIOCLIM variables 1-11. A. Scree p plot for BIOCLIM 1-11 (temperature). B. Loadings for PC1 (temperature). C. Loadings for PC2 (temperature). D. Scree plot for BIOCLIM 12-19 (precipitation). E. Loadings for PC1 (precipitation). F. Loadings for PC2 (precipitation).
