## Supplementary data 1-7 for "Genetic, morphological, and niche variation in the widely hybridizing *Rhus integrifolia-Rhus ovata* species complex": S7.docx

**Table S7**. Eigenvalues and variance components for PCA of environmental variables for GBIF herbarium records (left) and collection localities in the current study (right).

|  | **Expanded**  **Data (GBIF)** |  | **Collection**  **localities (n = 19)** |  |
| --- | --- | --- | --- | --- |
| **PC** | **Eigenvalue** | **% variance** | **Eigenvalue** | **% variance** |
| 1 | 7.9563 | 49.727 | 8.46291 | 44.542 |
| 2 | 1.68063 | 10.504 | 5.46527 | 28.765 |
| 3 | 1.21356 | 7.5848 | 2.42125 | 12.743 |
| 4 | 1.11714 | 6.9821 | 1.30401 | 6.8632 |
| 5 | 0.958839 | 5.9927 | 0.647778 | 3.4094 |
| 6 | 0.712245 | 4.4515 | 0.399497 | 2.1026 |
| 7 | 0.640414 | 4.0026 | 0.138317 | 0.72799 |
| 8 | 0.464707 | 2.9044 | 0.080852 | 0.42554 |
| 9 | 0.373972 | 2.3373 | 0.047957 | 0.25241 |
| 10 | 0.307387 | 1.9212 | 0.019478 | 0.10252 |
| 11 | 0.238572 | 1.4911 | 0.005659 | 0.029782 |
| 12 | 0.212436 | 1.3277 | 0.003171 | 0.016687 |
| 13 | 0.075412 | 0.47132 | 0.002359 | 0.012417 |
| 14 | 0.047726 | 0.29829 | 0.000816 | 0.004294 |
| 15 | 0.000478 | 0.002987 | 0.000345 | 0.001814 |
| 16 | 0.000178 | 0.001115 | 0.000233 | 0.001228 |
