## Supplementary data 8 for "Genetic, morphological, and niche variation in the widely hybridizing *Rhus integrifolia-Rhus ovata* species complex": maxent.html

Maxent model

### Maxent model

  
 This page contains some analysis of the Maxent model result, created Wed Jan 08 19:50:12 EST 2020 using 'dismo' version 1.1-4 & Maxent version 3.4.1. If you would like to do further analyses, the raw data used here is linked to at the end of this page.  
  

---

#### Analysis of omission/commission

The following picture shows the omission rate and predicted area as a function of the cumulative threshold. The omission rate is is calculated both on the training presence records, and (if test data are used) on the test records. The omission rate should be close to the predicted omission, because of the definition of the cumulative threshold.
  
  
  
 The next picture is the receiver operating characteristic (ROC) curve for the same data. Note that the specificity is defined using predicted area, rather than true commission (see the paper by Phillips, Anderson and Schapire cited on the help page for discussion of what this means). This implies that the maximum achievable AUC is less than 1. If test data is drawn from the Maxent distribution itself, then the maximum possible test AUC would be 0.983 rather than 1; in practice the test AUC may exceed this bound.
  
  
  
  
Some common thresholds and corresponding omission rates are as follows. If test data are available, binomial probabilities are calculated exactly if the number of test samples is at most 25, otherwise using a normal approximation to the binomial. These are 1-sided p-values for the null hypothesis that test points are predicted no better than by a random prediction with the same fractional predicted area. The "Balance" threshold minimizes 6 \* training omission rate + .04 \* cumulative threshold + 1.6 \* fractional predicted area.  
  

|  |  |  |  |  |  |  |  |  |  |  |  |  |  |  |  |  |  |  |  |  |  |  |  |  |  |  |  |  |  |  |  |  |  |  |  |  |  |  |  |  |  |  |  |  |  |  |  |  |  |
| --- | --- | --- | --- | --- | --- | --- | --- | --- | --- | --- | --- | --- | --- | --- | --- | --- | --- | --- | --- | --- | --- | --- | --- | --- | --- | --- | --- | --- | --- | --- | --- | --- | --- | --- | --- | --- | --- | --- | --- | --- | --- | --- | --- | --- | --- | --- | --- | --- | --- |
| Cumulative threshold | Cloglog threshold | Description | Fractional predicted area | Training omission rate || 1.000 | 0.006 | Fixed cumulative value 1 | 0.082 | 0.000 || 5.000 | 0.125 | Fixed cumulative value 5 | 0.039 | 0.032 || 10.000 | 0.346 | Fixed cumulative value 10 | 0.030 | 0.061 || 1.220 | 0.012 | Minimum training presence | 0.071 | 0.000 || 12.261 | 0.416 | 10 percentile training presence | 0.028 | 0.097 || 7.576 | 0.207 | Equal training sensitivity and specificity | 0.033 | 0.032 || 4.370 | 0.107 | Maximum training sensitivity plus specificity | 0.042 | 0.022 || 1.220 | 0.012 | Balance training omission, predicted area and threshold value | 0.071 | 0.000 || 4.344 | 0.106 | Equate entropy of thresholded and original distributions | 0.042 | 0.022 |

  
  
(A link to the Explain tool was not made for this model. The model uses product features, while the Explain tool can only be used for additive models.)  
  
  

---

#### Response curves

  
These curves show how each environmental variable affects the Maxent prediction.
The
curves show how the predicted probability of presence changes as each environmental variable is varied, keeping all other environmental variables at their average sample value. Click on a response curve to see a larger version. Note that the curves can be hard to interpret if you have strongly correlated variables, as the model may depend on the correlations in ways that are not evident in the curves. In other words, the curves show the marginal effect of changing exactly one variable, whereas the model may take advantage of sets of variables changing together.  
  
 
 
 
 
 
 
 
 
 
 
  
  
In contrast to the above marginal response curves, each of the following curves represents a different model, namely, a Maxent model created using only the corresponding variable. These plots reflect the dependence of predicted suitability both on the selected variable and on dependencies induced by correlations between the selected variable and other variables. They may be easier to interpret if there are strong correlations between variables.  
  
 
 
 
 
 
 
 
 
 
 
  
  

---

#### Analysis of variable contributions

  
The following table gives estimates of relative contributions of the environmental variables to the Maxent model. To determine the first estimate, in each iteration of the training algorithm, the increase in regularized gain is added to the contribution of the corresponding variable, or subtracted from it if the change to the absolute value of lambda is negative. For the second estimate, for each environmental variable in turn, the values of that variable on training presence and background data are randomly permuted. The model is reevaluated on the permuted data, and the resulting drop in training AUC is shown in the table, normalized to percentages. As with the variable jackknife, variable contributions should be interpreted with caution when the predictor variables are correlated.  
  

|  |  |  |  |  |  |  |
| --- | --- | --- | --- | --- | --- | --- |
| Variable | Percent contribution | Permutation importance || X18 | 55.2 | 5.5 |
| X14 | 18.3 | 14.3 |
| X15 | 9.8 | 0.3 |
| X19 | 6.6 | 11.4 |
| X08 | 3.7 | 11.5 |
| X02 | 2.9 | 14.4 |
| X09 | 1.5 | 6.8 |
| X12 | 0.8 | 9.4 |
| X03 | 0.7 | 15.4 |
| X01.1 | 0.4 | 10.8 |

  
  
The following picture shows the results of the jackknife test of variable importance. The environmental variable with highest gain when used in isolation is X18, which therefore appears to have the most useful information by itself. The environmental variable that decreases the gain the most when it is omitted is X02, which therefore appears to have the most information that isn't present in the other variables.  
  
  
  

---

#### Raw data outputs and control parameters

  
The data used in the above analysis is contained in the next links. Please see the Help button for more information on these.  
The model applied to the training environmental layers  
The coefficients of the model  
The omission and predicted area for varying cumulative and raw thresholds  
The prediction strength at the training and (optionally) test presence sites  
Results for all species modeled in the same Maxent run, with summary statistics and (optionally) jackknife results  
  
  
Regularized training gain is 3.177, training AUC is 0.985, unregularized training gain is 3.260.  
Algorithm terminated after 500 iterations (15 seconds).  
  
The follow settings were used during the run:  
278 presence records used for training.  
10278 points used to determine the Maxent distribution (background points and presence points).  
Environmental layers used (all continuous): X01.1 X02 X03 X08 X09 X12 X14 X15 X18 X19  
Regularization values: linear/quadratic/product: 0.050, categorical: 0.250, threshold: 1.000, hinge: 0.500  
Feature types used: hinge product linear quadratic  
responsecurves: true  
jackknife: true  
outputdirectory: Rhus integrifolia\_contemp\_Pearson  
samplesfile: Rhus integrifolia\_contemp\_Pearson/presence  
environmentallayers: Rhus integrifolia\_contemp\_Pearson/absence  
randomseed: true  
replicates: 0  
autorun: true  
visible: false  
threads: 28  
verbose: true  
Command line used: autorun -e Rhus integrifolia\_contemp\_Pearson/absence -o Rhus integrifolia\_contemp\_Pearson -s Rhus integrifolia\_contemp\_Pearson/presence -z -J replicates=0 randomseed=TRUE writeclampgrid=TRUE writemess=TRUE plots=TRUE threads=28 verbose=TRUE outputgrids=TRUE pictures=TRUE responsecurves=TRUE  
  
