## Supplementary data 8 for "Genetic, morphological, and niche variation in the widely hybridizing *Rhus integrifolia-Rhus ovata* species complex": maxent.html

Maxent model

### Maxent model

  
 This page contains some analysis of the Maxent model result, created Wed Jan 08 19:52:51 EST 2020 using 'dismo' version 1.1-4 & Maxent version 3.4.1. If you would like to do further analyses, the raw data used here is linked to at the end of this page.  
  

---

#### Analysis of omission/commission

The following picture shows the omission rate and predicted area as a function of the cumulative threshold. The omission rate is is calculated both on the training presence records, and (if test data are used) on the test records. The omission rate should be close to the predicted omission, because of the definition of the cumulative threshold.
  
  
  
 The next picture is the receiver operating characteristic (ROC) curve for the same data. Note that the specificity is defined using predicted area, rather than true commission (see the paper by Phillips, Anderson and Schapire cited on the help page for discussion of what this means). This implies that the maximum achievable AUC is less than 1. If test data is drawn from the Maxent distribution itself, then the maximum possible test AUC would be 0.966 rather than 1; in practice the test AUC may exceed this bound.
  
  
  
  
Some common thresholds and corresponding omission rates are as follows. If test data are available, binomial probabilities are calculated exactly if the number of test samples is at most 25, otherwise using a normal approximation to the binomial. These are 1-sided p-values for the null hypothesis that test points are predicted no better than by a random prediction with the same fractional predicted area. The "Balance" threshold minimizes 6 \* training omission rate + .04 \* cumulative threshold + 1.6 \* fractional predicted area.  
  

|  |  |  |  |  |  |  |  |  |  |  |  |  |  |  |  |  |  |  |  |  |  |  |  |  |  |  |  |  |  |  |  |  |  |  |  |  |  |  |  |  |  |  |  |  |  |  |  |  |  |
| --- | --- | --- | --- | --- | --- | --- | --- | --- | --- | --- | --- | --- | --- | --- | --- | --- | --- | --- | --- | --- | --- | --- | --- | --- | --- | --- | --- | --- | --- | --- | --- | --- | --- | --- | --- | --- | --- | --- | --- | --- | --- | --- | --- | --- | --- | --- | --- | --- | --- |
| Cumulative threshold | Cloglog threshold | Description | Fractional predicted area | Training omission rate || 1.000 | 0.009 | Fixed cumulative value 1 | 0.177 | 0.000 || 5.000 | 0.124 | Fixed cumulative value 5 | 0.083 | 0.019 || 10.000 | 0.301 | Fixed cumulative value 10 | 0.064 | 0.049 || 1.532 | 0.019 | Minimum training presence | 0.142 | 0.000 || 15.042 | 0.447 | 10 percentile training presence | 0.055 | 0.099 || 11.653 | 0.358 | Equal training sensitivity and specificity | 0.060 | 0.060 || 4.282 | 0.096 | Maximum training sensitivity plus specificity | 0.089 | 0.010 || 1.532 | 0.019 | Balance training omission, predicted area and threshold value | 0.142 | 0.000 || 4.583 | 0.110 | Equate entropy of thresholded and original distributions | 0.086 | 0.018 |


|  |  |  |  |  |  |  |
| --- | --- | --- | --- | --- | --- | --- |
| Variable | Percent contribution | Permutation importance || X14 | 48.2 | 22.9 |
| X19 | 35 | 64.2 |
| X18 | 4.9 | 2.1 |
| X12 | 3.8 | 3.2 |
| X01.1 | 2.2 | 2.6 |
| X08 | 1.7 | 1 |
| X15 | 1.5 | 0.4 |
| X02 | 1.3 | 0.7 |
| X03 | 1.1 | 3 |
| X09 | 0.3 | 0.1 |

  
  
The following picture shows the results of the jackknife test of variable importance. The environmental variable with highest gain when used in isolation is X14, which therefore appears to have the most useful information by itself. The environmental variable that decreases the gain the most when it is omitted is X19, which therefore appears to have the most information that isn't present in the other variables.  
  
  
  

---

#### Raw data outputs and control parameters

  
The data used in the above analysis is contained in the next links. Please see the Help button for more information on these.  
The model applied to the training environmental layers  
The coefficients of the model  
The omission and predicted area for varying cumulative and raw thresholds  
The prediction strength at the training and (optionally) test presence sites  
Results for all species modeled in the same Maxent run, with summary statistics and (optionally) jackknife results  
  
  
Regularized training gain is 2.451, training AUC is 0.970, unregularized training gain is 2.567.  
Algorithm terminated after 500 iterations (15 seconds).  
  
The follow settings were used during the run:  
514 presence records used for training.  
10514 points used to determine the Maxent distribution (background points and presence points).  
Environmental layers used (all continuous): X01.1 X02 X03 X08 X09 X12 X14 X15 X18 X19  
Regularization values: linear/quadratic/product: 0.050, categorical: 0.250, threshold: 1.000, hinge: 0.500  
Feature types used: hinge product linear quadratic  
responsecurves: true  
jackknife: true  
outputdirectory: Rhus ovata\_contemp\_Pearson  
samplesfile: Rhus ovata\_contemp\_Pearson/presence  
environmentallayers: Rhus ovata\_contemp\_Pearson/absence  
randomseed: true  
replicates: 0  
autorun: true  
visible: false  
threads: 28  
verbose: true  
Command line used: autorun -e Rhus ovata\_contemp\_Pearson/absence -o Rhus ovata\_contemp\_Pearson -s Rhus ovata\_contemp\_Pearson/presence -z -J replicates=0 randomseed=TRUE writeclampgrid=TRUE writemess=TRUE plots=TRUE threads=28 verbose=TRUE outputgrids=TRUE pictures=TRUE responsecurves=TRUE  
  
