## Supplementary figures and images for "Genetic, morphological, and niche variation in the widely hybridizing *Rhus integrifolia-Rhus ovata* species complex"

### species_jacknife.png

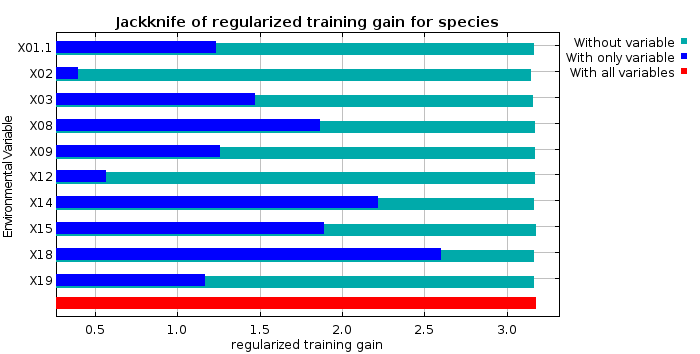

### species_jacknife.png

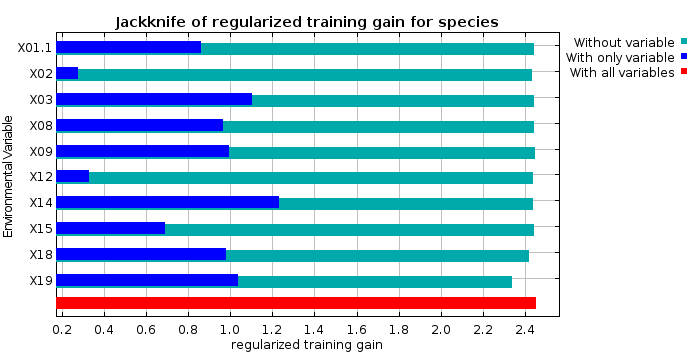

### species_omission.png

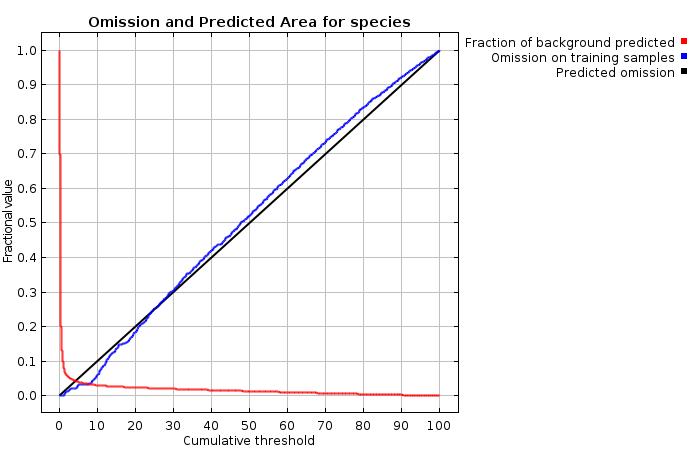

### species_omission.png

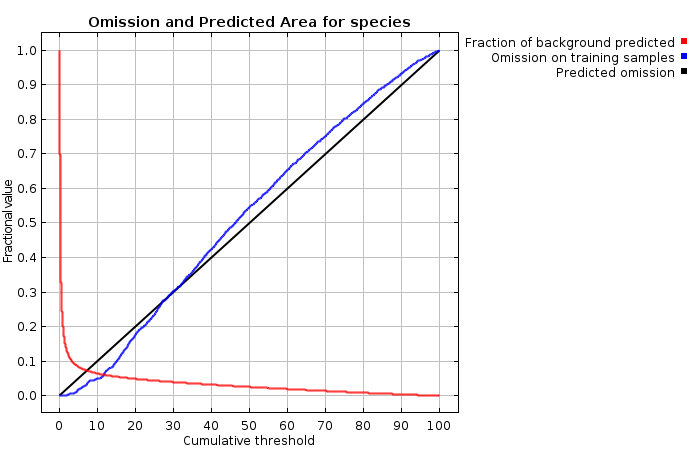

### species_roc.png

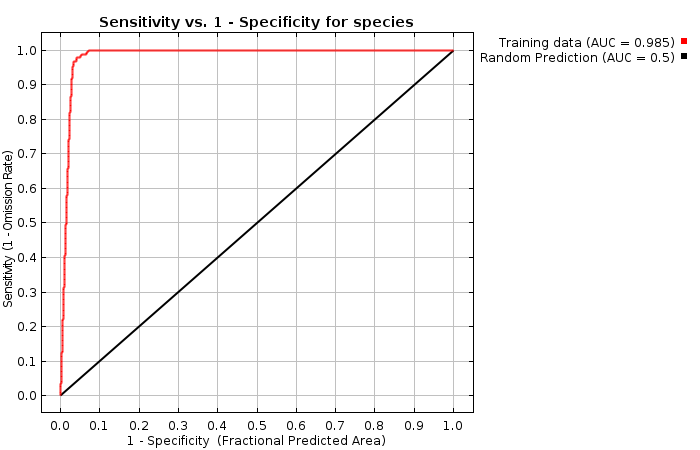

### species_roc.png

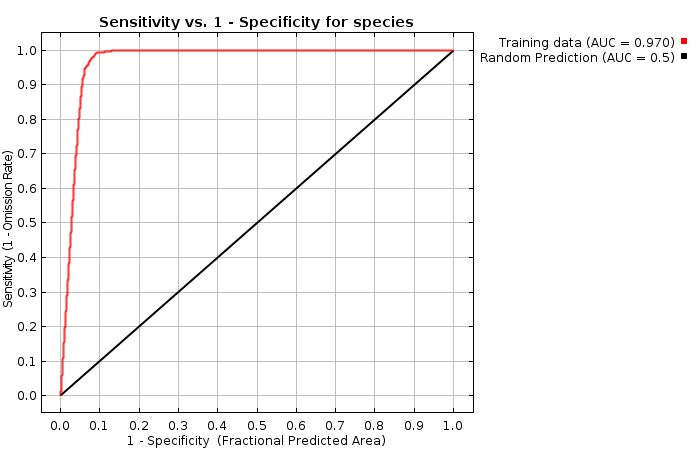

### species_X01.1.png

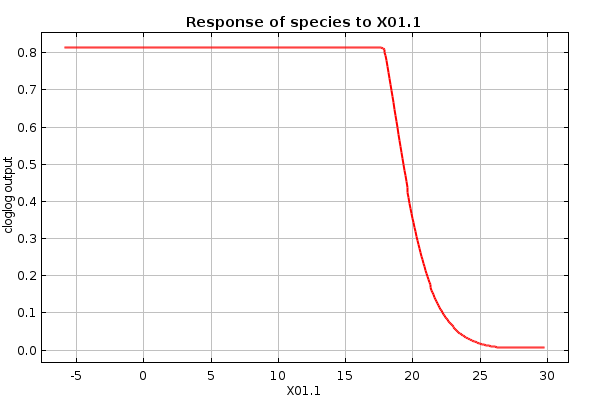

### species_X01.1.png

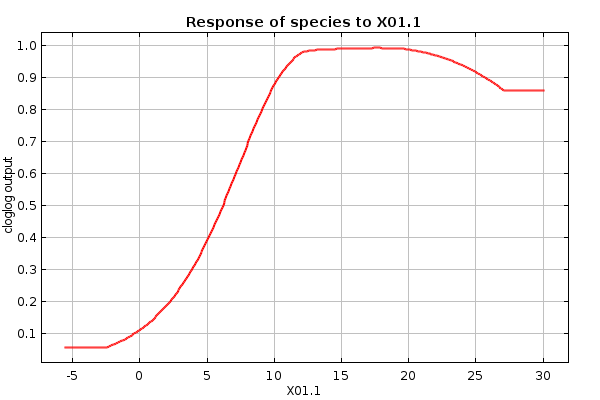

### species_X01.1_only.png

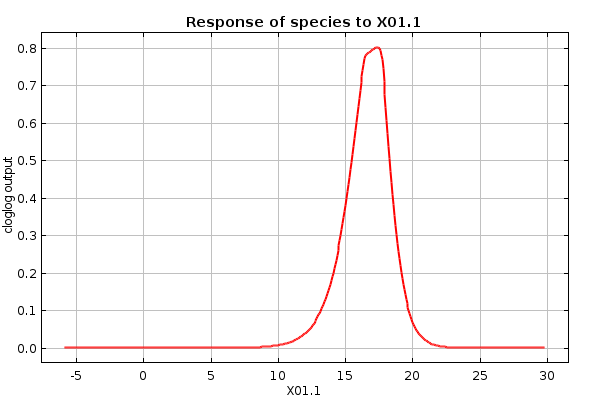

### species_X01.1_only.png

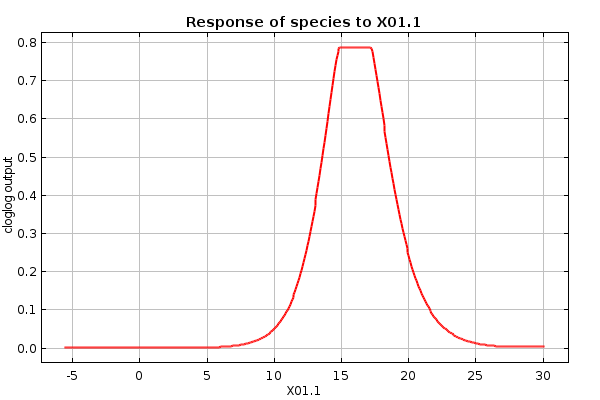

### species_X01.1_only_thumb.png

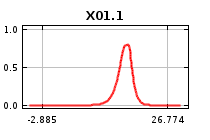

### species_X01.1_only_thumb.png

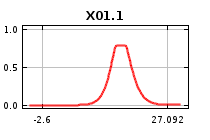

### species_X01.1_thumb.png

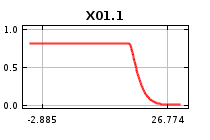

### species_X01.1_thumb.png

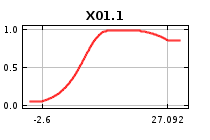

### species_X02.png

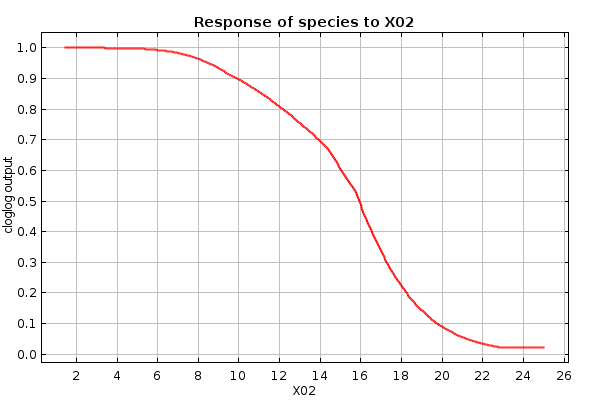

### species_X02.png

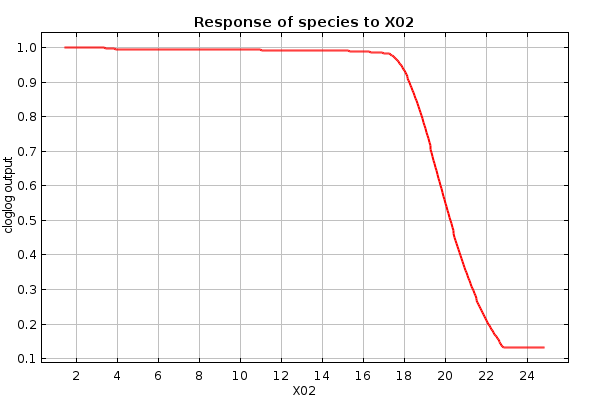

### species_X02_only.png

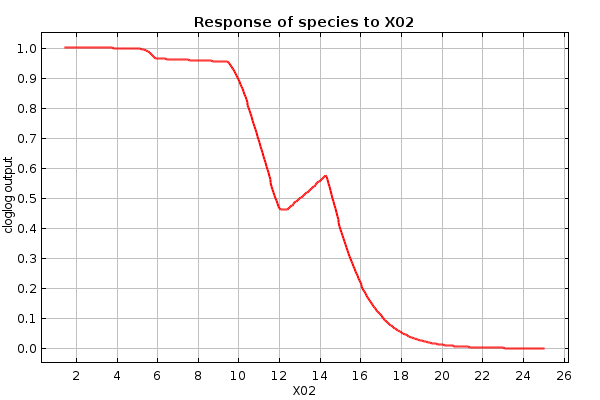

### species_X02_only.png

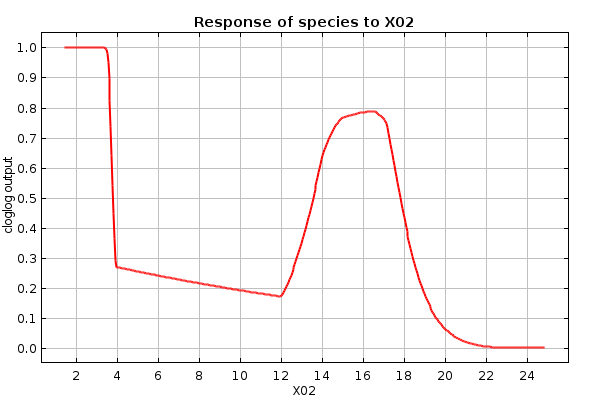

### species_X02_only_thumb.png

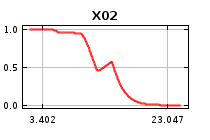

### species_X02_only_thumb.png

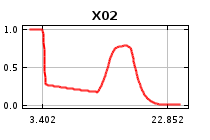

### species_X02_thumb.png

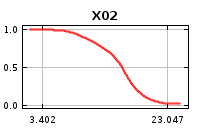

### species_X02_thumb.png

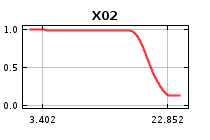

### species_X03.png

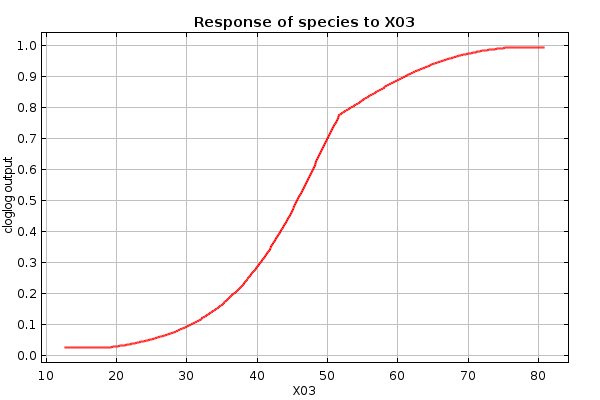

### species_X03.png

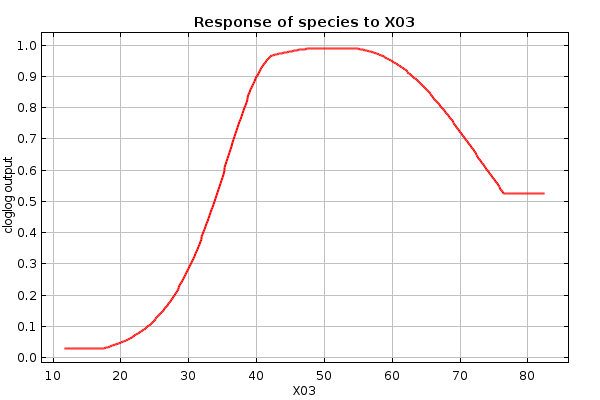

### species_X03_only.png

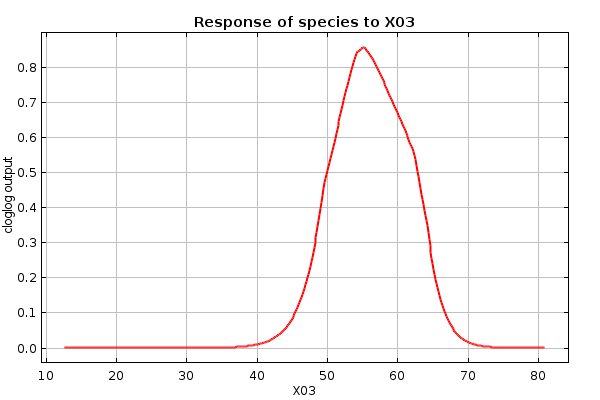

### species_X03_only.png

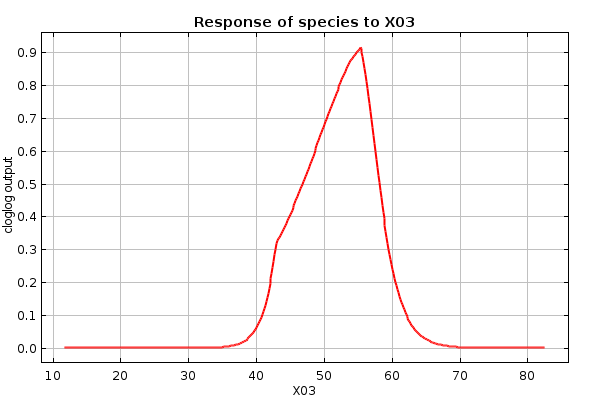

### species_X03_only_thumb.png

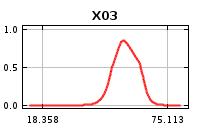

### species_X03_only_thumb.png

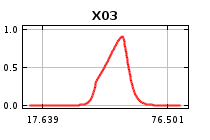

### species_X03_thumb.png

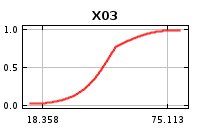

### species_X03_thumb.png

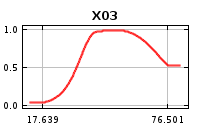

### species_X08.png

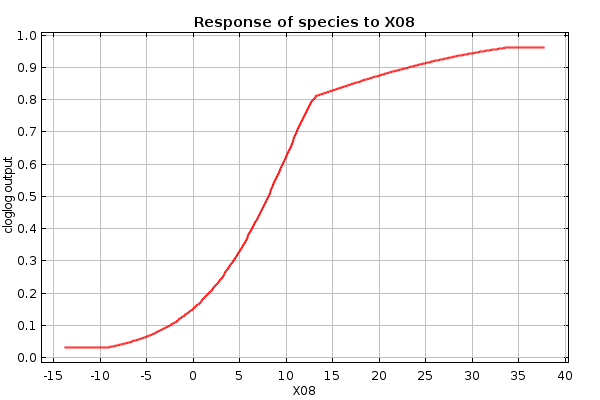

### species_X08.png

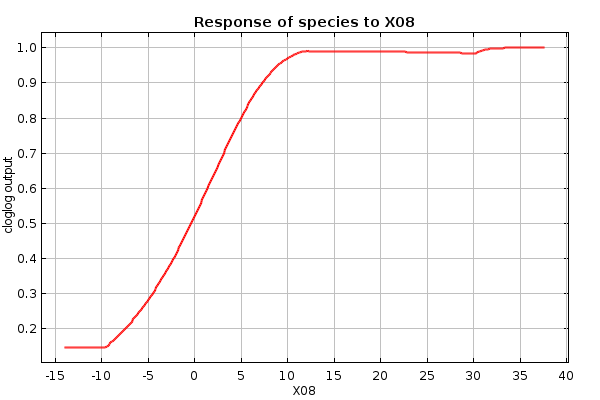

### species_X08_only.png

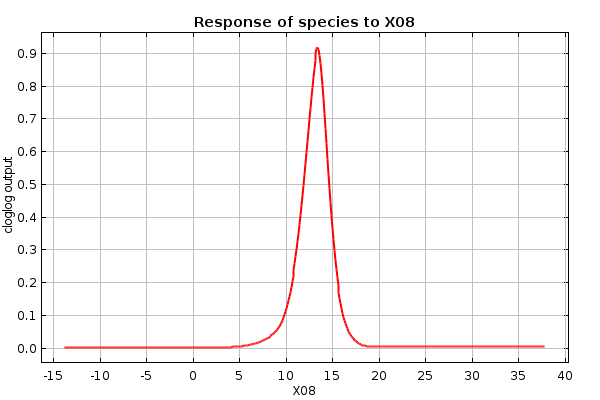

### species_X08_only.png

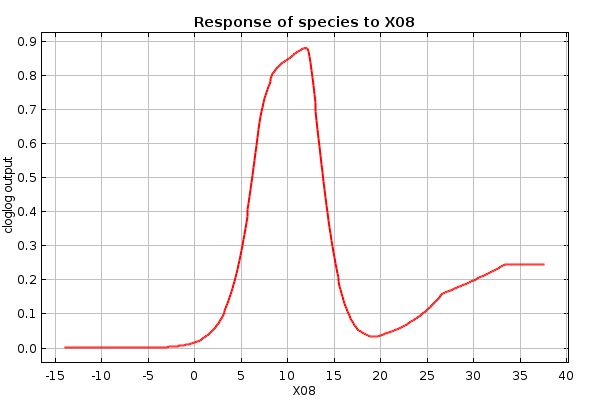

### species_X08_only_thumb.png

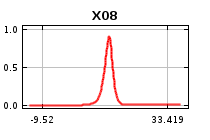

### species_X08_only_thumb.png

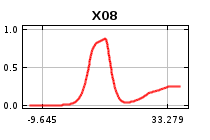

### species_X08_thumb.png

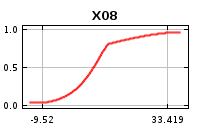

### species_X08_thumb.png

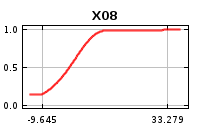

### species_X09.png

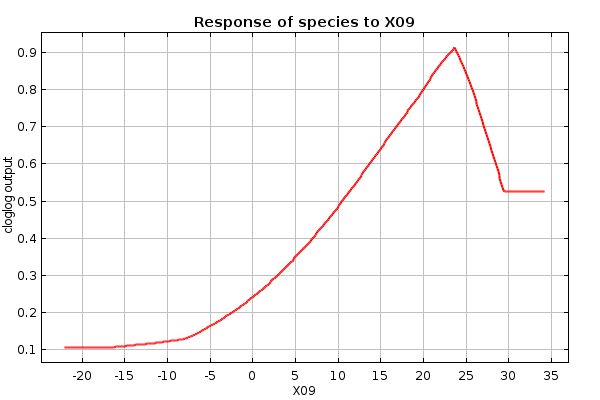

### species_X09.png

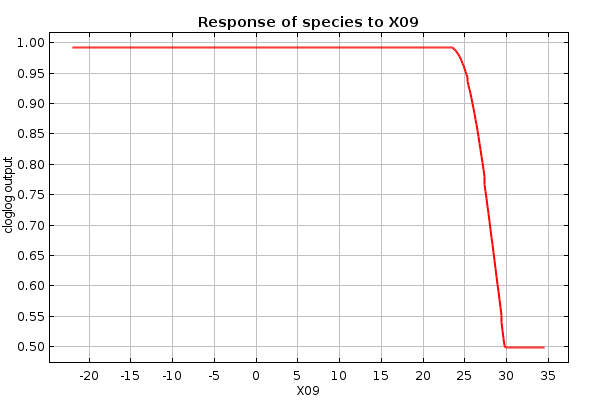

### species_X09_only.png

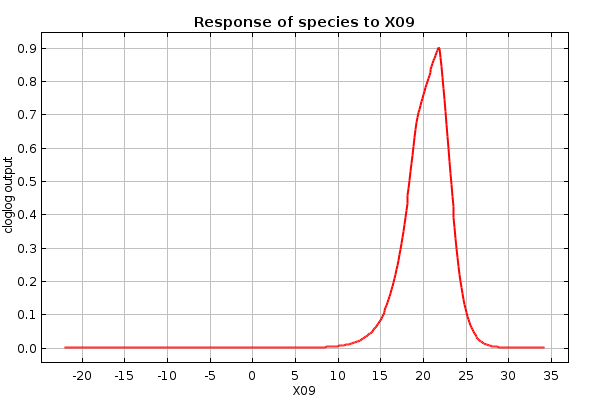

### species_X09_only.png

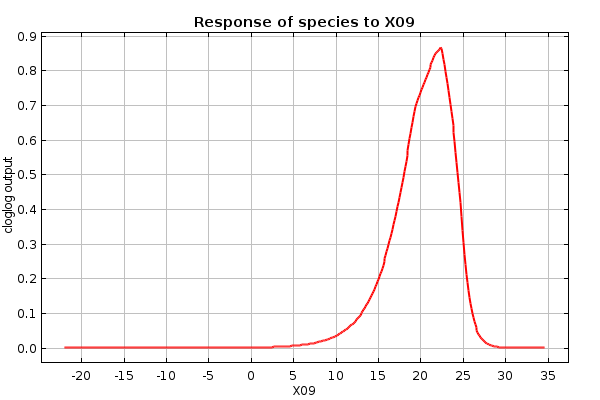

### species_X09_only_thumb.png

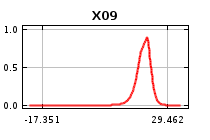

### species_X09_only_thumb.png

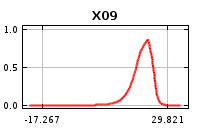

### species_X09_thumb.png

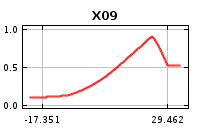

### species_X09_thumb.png

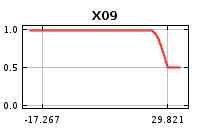

### species_X12.png

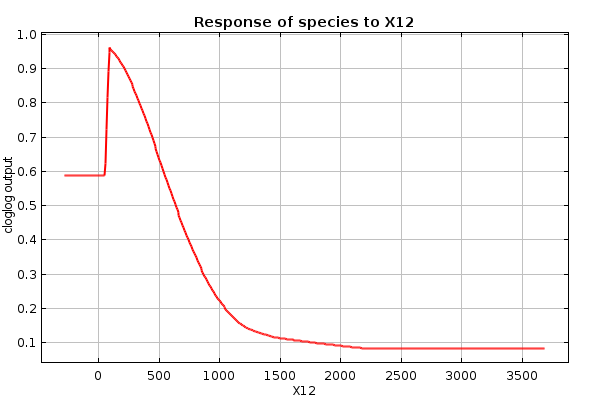

### species_X12.png

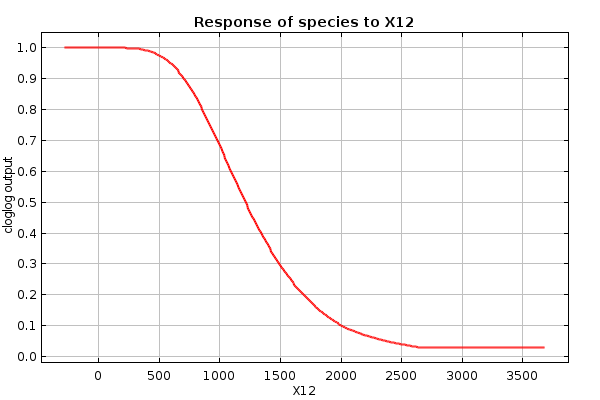

### species_X12_only.png

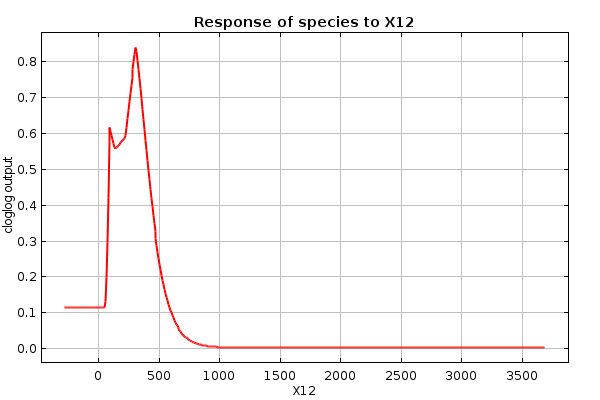

### species_X12_only.png

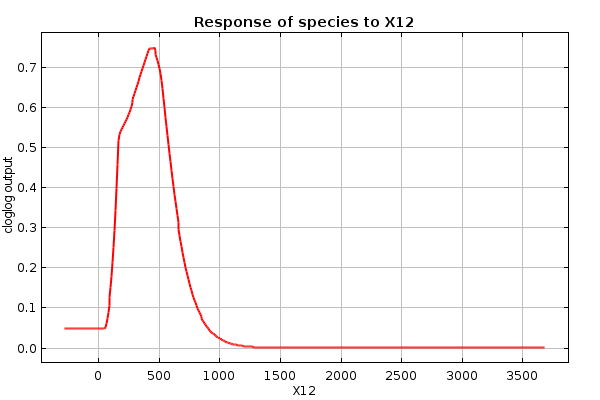

### species_X12_only_thumb.png

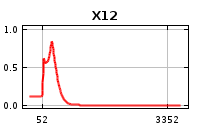

### species_X12_only_thumb.png

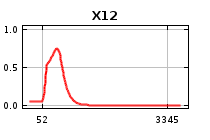

### species_X12_thumb.png

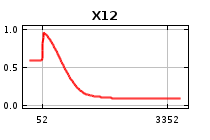

### species_X12_thumb.png

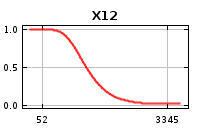

### species_X14.png

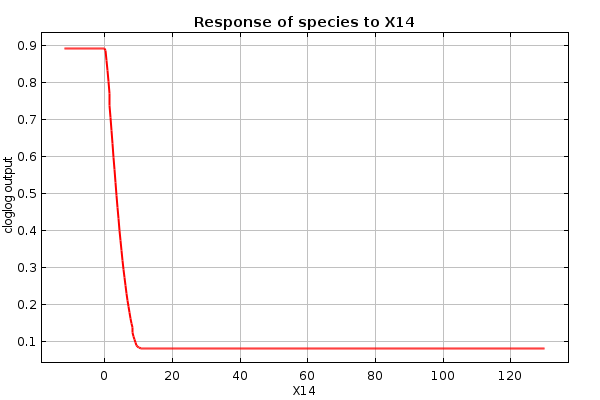

### species_X14.png

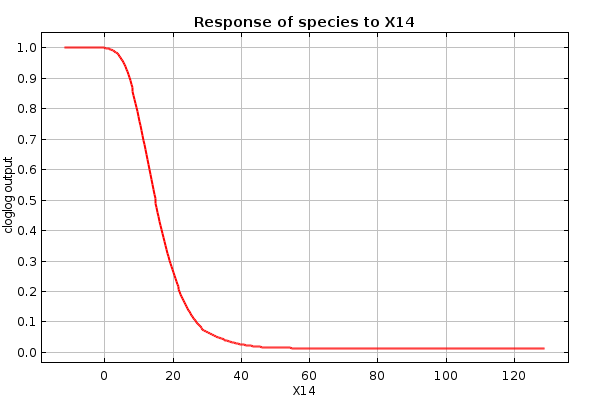

### species_X14_only.png

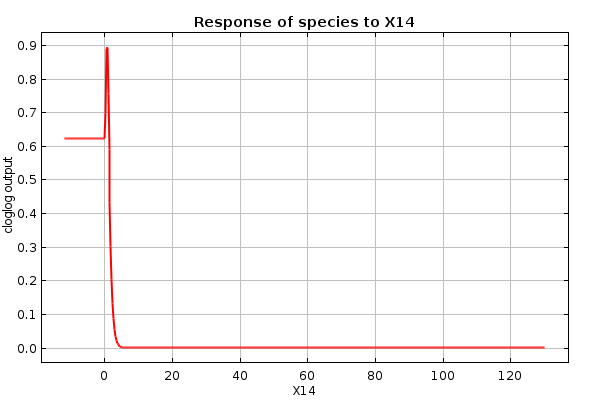

### species_X14_only.png

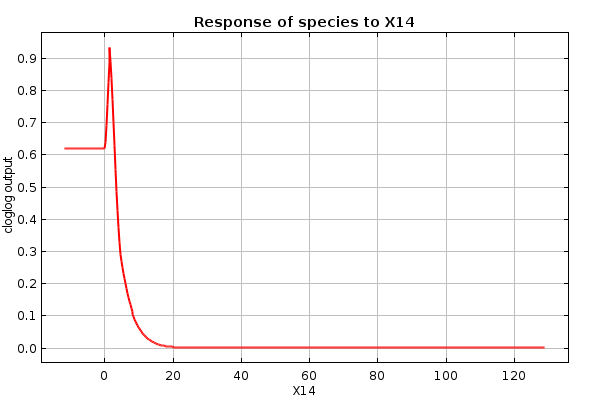

### species_X14_only_thumb.png

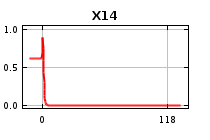

### species_X14_only_thumb.png

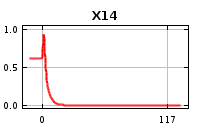

### species_X14_thumb.png

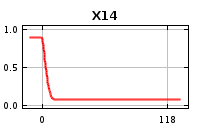

### species_X14_thumb.png

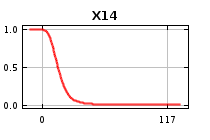

### species_X15.png

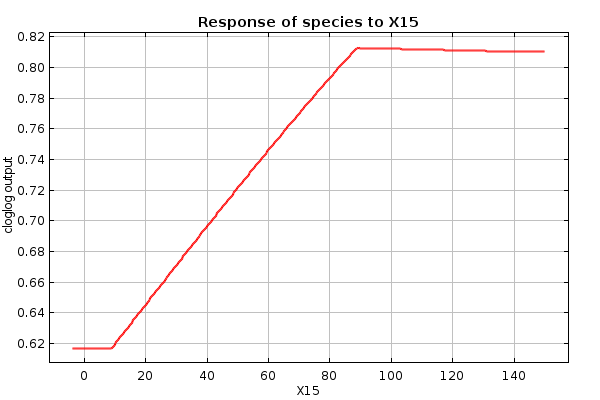

### species_X15.png

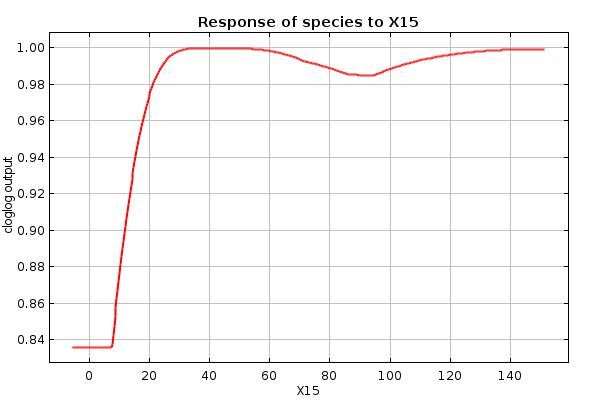

### species_X15_only.png

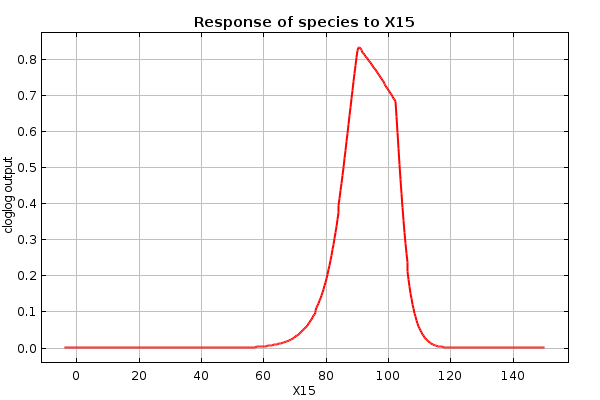

### species_X15_only.png

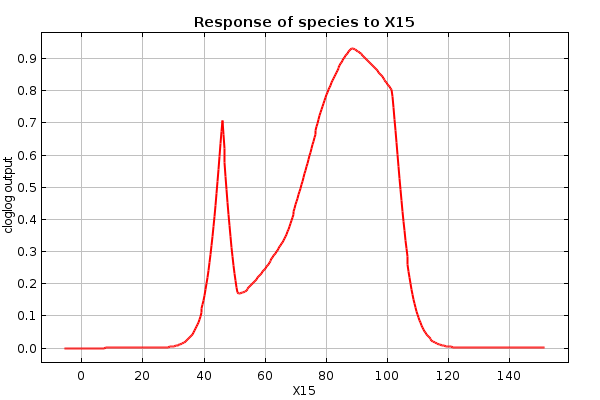

### species_X15_only_thumb.png

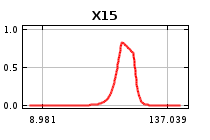

### species_X15_only_thumb.png

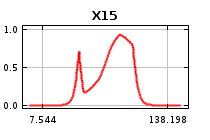

### species_X15_thumb.png

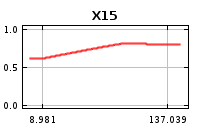

### species_X15_thumb.png

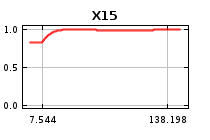

### species_X18.png

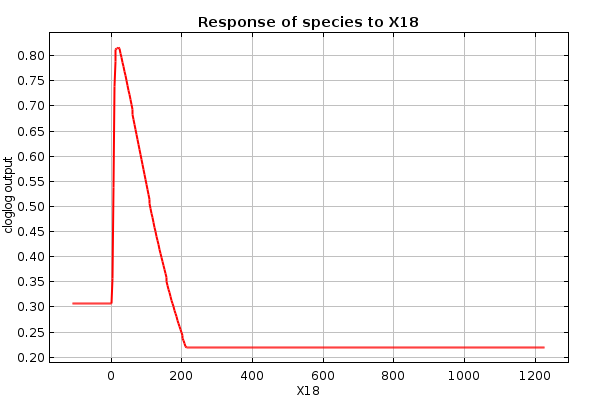

### species_X18.png

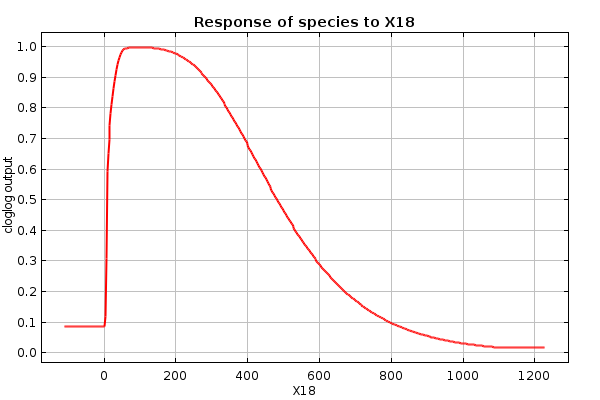

### species_X18_only.png

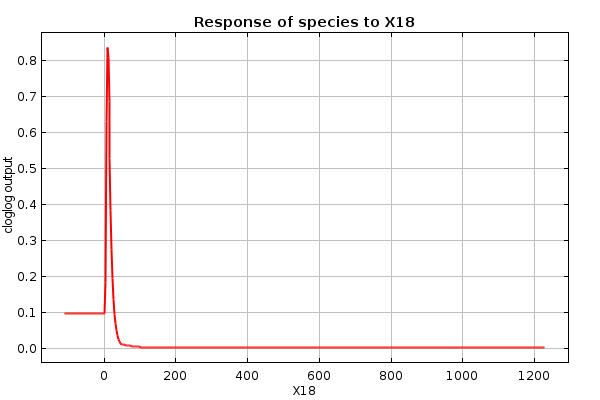

### species_X18_only.png

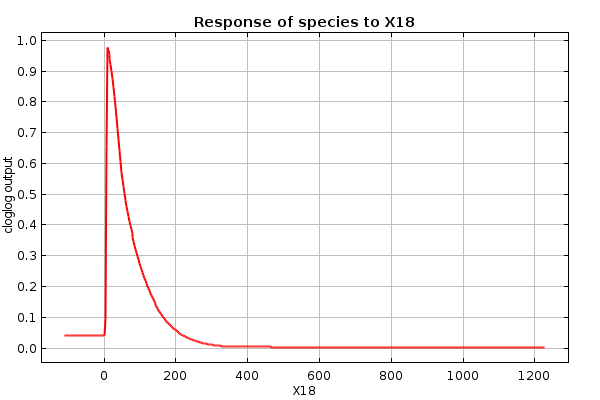

### species_X18_only_thumb.png

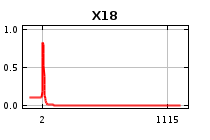

### species_X18_only_thumb.png

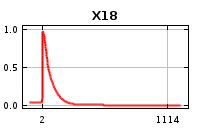

### species_X18_thumb.png

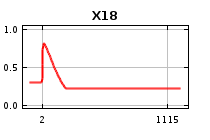

### species_X18_thumb.png

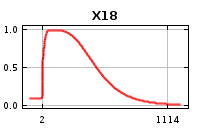

### species_X19.png

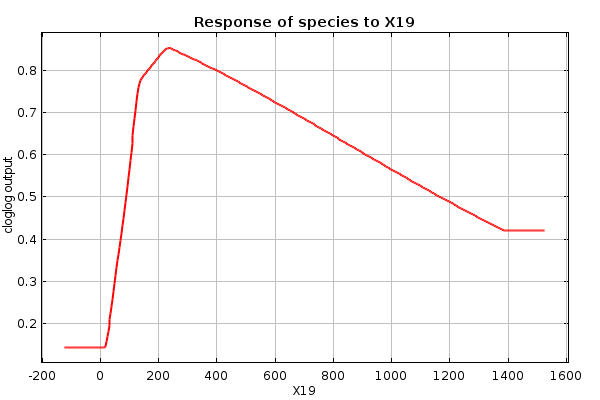

### species_X19.png

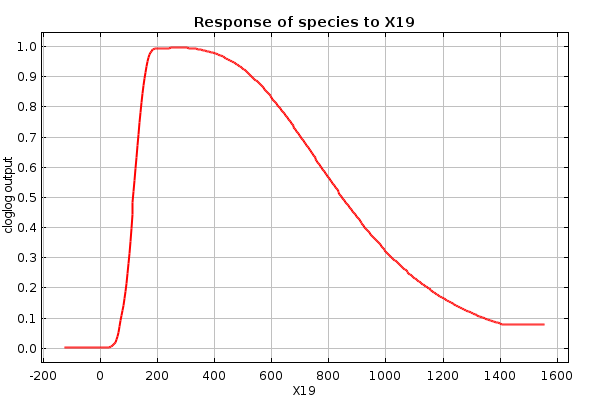

### species_X19_only.png

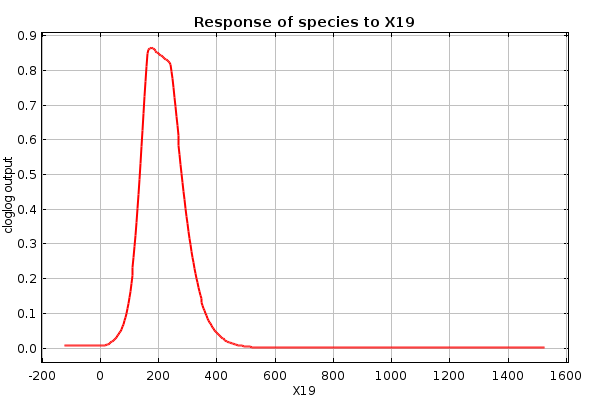

### species_X19_only.png

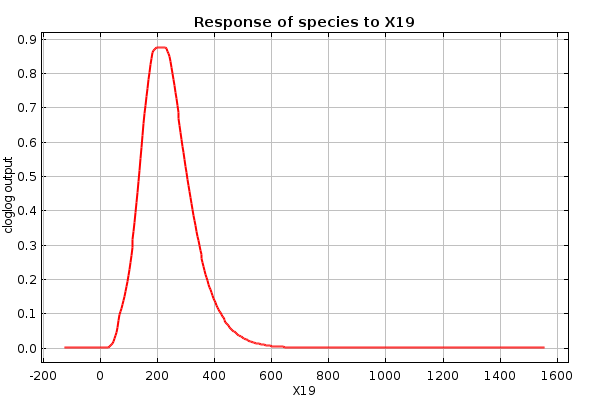

### species_X19_only_thumb.png

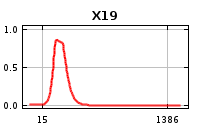

### species_X19_only_thumb.png

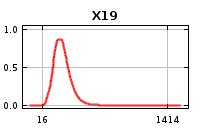

### species_X19_thumb.png

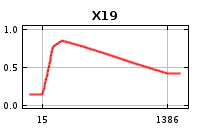

### species_X19_thumb.png

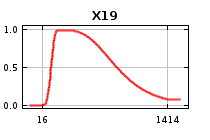
